# A single-cell expression simulator guided by gene regulatory networks

**DOI:** 10.1101/716811

**Authors:** Payam Dibaeinia, Saurabh Sinha

**Affiliations:** Department of Computer Science, University of Illinois Urbana-Champaign, Urbana, IL, 61801, USA; Carl R. Woese Institute of Genomic Biology, University of Illinois Urbana-Champaign, Urbana, IL, 61801, USA; Cancer Center at Illinois, University of Illinois Urbana-Champaign, Urbana, IL, 61801, USA

## Abstract

A common approach to benchmarking of single-cell transcriptomics tools is to generate synthetic data sets that resemble experimental data in their statistical properties. However, existing single-cell simulators do not incorporate known principles of transcription factor-gene regulatory interactions that underlie expression dynamics. Here we present SERGIO, a simulator of single-cell gene expression data that models the stochastic nature of transcription as well as linear and non-linear influences of multiple transcription factors on genes according to a user-provided gene regulatory network. SERGIO is capable of simulating any number of cell types in steady-state or cells differentiating to multiple fates according to a provided trajectory, reporting both unspliced and spliced transcript counts in single-cells. We show that data sets generated by SERGIO are comparable with experimental data in terms of multiple statistical measures. We also illustrate the use of SERGIO to benchmark several popular single-cell analysis tools, including GRN inference methods.

## Introduction

Single-cell transcriptomics technologies are revolutionizing biology today^1–4^, and have led to the rapid development of computational tools for analyzing the resulting data sets^5–8^. These tools, developed for a wide array of tasks such as clustering^9–11^, trajectory inference^12,13^ and gene regulatory network (GRN) reconstruction^9,14,15^, as well as pre-processing operations such as imputation^16–18^, adopt complementary strategies whose relative merits and weaknesses are not clear *a priori*. In some cases, single-cell data sets annotated using domain knowledge^19,20^ allow objective evaluations of different strategies, but this is not a scalable approach to systematic benchmarking. A promising alternative approach is to synthesize single-cell expression data sets that mimic real data in their statistical properties and for which underlying biological relationships are known by construction.

Simulation tools (“simulators”) for single-cell expression data have been reported in various forms. Several studies offering novel analysis tools use in-house simulators to benchmark those tools^8,21–26^, while other studies specifically develop simulators for use by the community^27–32^. Most of these simulators are geared towards capturing the noise characteristics of technologies such as single-cell RNA-seq (scRNA-seq), by first estimating statistical quantities describing real data sets and then sampling single-cell expression profiles from probability distributions that mirror those quantities. A crucial aspect of biology missing in current simulators is the gene regulatory network (GRN): the set of transcription factor (TF)-gene relationships that underlies the dynamics and steady states of gene expression in each cell. We believe it is imperative that a singlecell expression simulator be guided by an underlying GRN, not only because of the biological realism that it represents, but also because this is the only direct way to benchmark tools specifically designed for GRN reconstruction. Some existing tools do attempt to induce gene-gene relationships in synthetic data using multi-gene statistical models for sampling purposes^28,33^, but these attempts do not incorporate the special properties of gene regulatory processes that have been reported in the literature^34–37^, including non-linear response to TFs, intrinsic fluctuations in expression and propagation of such “biological noise” along the GRN.

In the realm of “bulk” transcriptomics GRN-driven simulations are already the norm, as exemplified by the simulation tool called GeneNetWeaver (GNW)^38^, which was used in a community-wide effort to benchmark numerous GRN reconstruction tools^39–42^. GNW is not meant to simulate scRNA-seq data, and though some studies have employed workarounds to use it for this purpose^14,43^, it is believed that such synthetic data do not exhibit the statistical characteristics of contemporary single-cell data sets^43^. Furthermore, such workarounds do not offer key features necessary for a single cell expression simulator, such as simulation of multiple cell types and cells differentiating from one cell type to another.

In this work, we develop a simulator tool that (1) uses a principled mathematical description of transcriptional regulatory processes to synthesize single-cell expression data associated with a specified GRN, (2) includes stochasticity of gene expression as an integral part of the process, thus capturing biological noise expected to manifest in cell-to-cell variability, and (3) incorporates various types of measurement errors (“technical noise”) that are typical of single-cell technologies. The new tool, called SERGIO (**S**ingle-cell **E**xp**R**ession of **G**enes **I**n silic**o**), is freely available as a stand-alone software package. It borrows some of its modeling assumptions from the widely used GNW simulator, but relinquishes the more complex features of GNW, such as a thermodynamics-based model of regulation and explicit modeling of translation processes, which would have necessitated use of poorly-understood parameters during simulation and slowed down simulations of large GRNs.

SERGIO uses a stochastic differential equation (SDE) called the chemical Langevin equation^44^ to simulate a gene’s expression dynamics as a function of the changing (or fluctuating) levels of its regulators (TFs), as prescribed by a fixed GRN. It performs such simulations for any pre-specified number of genes in parallel, and generates single-cell expression “profiles” (expression values of all genes) by sampling from these temporal simulations in steady-state, thus mimicking established cell types. It allows users to specify the number of cell types to be simulated, via steady-state levels of a few “master” regulators in the GRN. SERGIO also allows users to simulate single-cell expression data from a specified differentiation program, for which it samples cells from transient portions of temporal simulations. In this simulation mode, SERGIO explicitly models the splicing step with an additional SDE, resulting in simulations of unspliced and spliced transcript levels. SERGIO subjects the synthesized expression data to a multi-step transformation where technical noise is incorporated in a manner reflecting real scRNA-seq data. To our knowledge, SERGIO is the first stand-alone simulator tool for single-cell transcriptomics that offers all of the above-mentioned features while basing its simulations on a given GRN. Here, we outline key aspects of its model and implementation and show that it may be used to generate realistic data sets that resemble an experimental scRNA-seq data set by several statistical measures. We then showcase its use to benchmark a number of popular single-cell analysis tools. We find that while modern tools are able to accurately identify cell types and differentiation trajectories from suitable data sets, their ability to reconstruct gene regulatory relationships remains severely limited.

## Results

We developed SERGIO to simulate how expression values of a specified number of genes vary from cell to cell under the control of a given GRN, and how such information is captured in modern single-cell RNA-seq data sets. We first simulate “clean” gene expression data based on the GRN and mathematical models of transcriptional processes, including stochasticity of such processes (“biological noise”). We then add “technical noise” to the clean data, mimicking the nature of measurement errors attributed to scRNA-seq technology^45^.

### Simulation of “clean” data

We generate expression profiles of single cells by sampling them from the steady state of a dynamical process that involves genes expressing at rates influenced by other genes (transcription factors) (Figure 1). A select few of the genes are pre-designated as master regulators (MRs); these have no regulatory inputs in the GRN and their expression evolves over time under constant production and decay rates (see Methods). Expression of every other gene (non-MR) evolves under a production rate determined by adding contributions from its GRN-specified regulators (equation 5 in Methods) and a constant decay rate. Each regulator’s contribution to a gene depends on the former’s current concentration and an interaction parameter (strength of activation or repression) specific to the regulator and regulated gene. This dependence is described by a Hill function^46^, thus allowing for non-linear effects.

**Figure 1:**
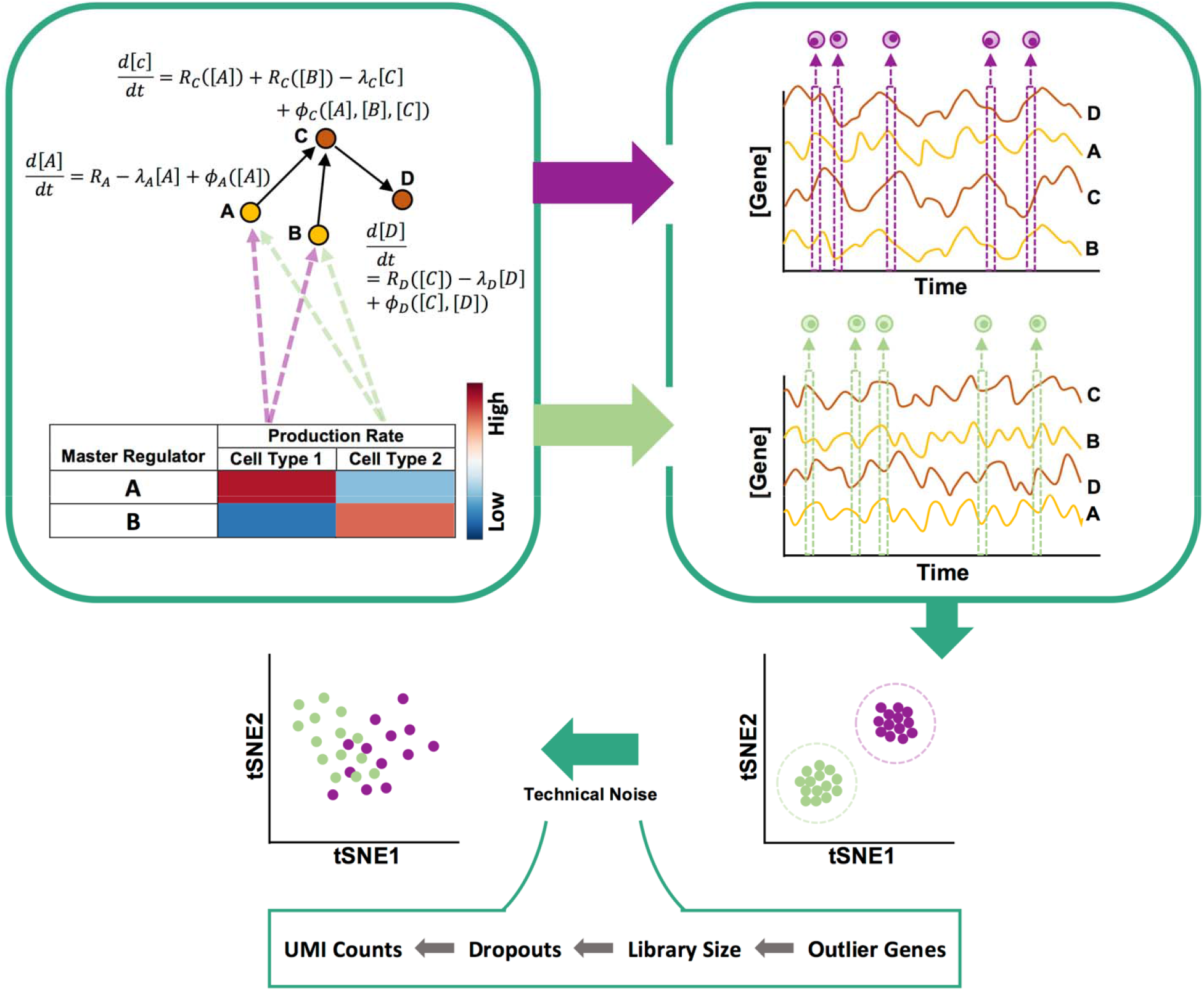
Overview of steady-state simulation pipeline. SERGIO uses stochastic differential equations (SDE) to describe the dynamics of mRNA transcripts of each gene (A,B,C,D) in a specified GRN (top left). Each gene’s SDE consists of a production rate which is modeled as the sum of contributions the gene receives from its regulators (e.g., from A and B for gene C). Such a contribution is modeled as a regulatory function (*R_gene_*) of the concentration of the TF except for “master regulators” (genes without regulators) for which the production rate is a constant (e.g., *R_A_*). Also, each SDE contains a term representing the decay of mRNA transcripts (e.g. *λ_C_[C]*) and a term representing biological noise (e.g., *ϕ_C_([A], [B], [C])*). A cell type is specified by the production rates of MRs, and SERGIO performs separate simulations for each cell type using these MR production rates. It initializes the concentration of genes to their estimated averaged steady-state concentrations and continues simulations in steady-state region for all genes simultaneously to generate time-course expression data (right). Finally, it samples singlecells from the time-course uniformly at random over the steady-state region. Bottom right: A cartoon illustration of the “clean” data generated by SERGIO shows single-cells tightly clustered by cell type. Upon adding technical noise, cells of different types become less well-separated but are still distinguishable by clustering algorithms.

Each gene’s time course is simulated while incorporating biological noise, using the chemical Langevin equation^44^, as adopted in the GeneNetWeaver (GNW) simulator^38^. Once the system of evolving expression profiles reaches steady state, we sample profiles from randomly selected time points. Variation in expression profiles across cells of the same type is assumed to mimic variation across time points in the steady state (the “ergodic assumption”^47^), hence the temporally sampled cells are used as the collection of cells in the synthetic data.

Specifying the fixed production rates of MRs determines the average steady state expression profile of the sampled cells, and is used to generate data for a single cell type. In order to synthesize a data set with multiple cell types, the above simulation is repeated using different settings of MR production rates. The aggregate of expression profiles sampled across all simulations forms the “clean” synthetic data set.

### Incorporation of technical noise

We then use the clean data to simulate integer-valued “count” data, as are produced in current scRNA-seq technologies, by sampling from a Poisson distribution whose mean is the real-valued expression level. However, prior to this conversion, the real-valued expression data matrix (genes × cells) is operated upon by modules that incorporate three different types of technical noise. The statistical details of these modules are borrowed from the Splatter simulation tool^32^ and re-implemented in SERGIO (see Methods).

### SERGIO simulates realistic data sets

We used SERGIO to generate synthetic data sets under three different settings of the underlying GRN, referred to as “data set 1” (DS1), “data set 2” (DS2) and “data set 3” (DS3). These three settings use GRNs with 100, 400 and 1200 genes respectively, that were sampled from real regulatory networks in *E. coli* or *S. cerevisae* (Table 1); all simulations included 300 cells for each of 9 cell types, for a total of 2700 single cells. Each data set was synthesized in 15 “replicates” by re-executing SERGIO with identical parameters multiple times. We sought to compare statistical properties of these synthetic data sets to a published data set from mouse brain comprising expression profiles of cells that are categorized into nine cell types with high confidence^48^, henceforth called the “real data set”. We thus configured SERGIO to introduce technical noise in the simulated expression profiles, to an extent that matches the real data set. This was done through manual iteration of the technical noise parameters (see Methods). For each simulation setting we sampled a comparison data set from the real data to have the same number of genes, repeating this 50 times to obtain 50 replicates of the (sampled) real data set, each of which was compared to the 15 replicates of the corresponding synthetic data set. We performed our comparisons using synthetic data with and without technical noise, referred to as the “noisy” and “clean” forms of the data set.

**Table 1:** Description of the synthetic data sets used in this study

| **Data Set ID** | **Network ID** | **Species** | **#Genes** | **#Regulators** | **#Edges** | **#Cell types** | **Differentiation** |
| --- | --- | --- | --- | --- | --- | --- | --- |
| DS1 | 2 | Ecoli | 100 | 10 | 258 | 9 | No |
| DS2 | 3 | Yeast | 400 | 37 | 1155 | 9 | No |
| DS3 | 4 | Ecoli | 1200 | 127 | 2713 | 9 | No |
| DS4 | 1 | Ecoli | 100 | 10 | 137 | 3 | Yes |
| DS5 | 1 | Ecoli | 100 | 10 | 137 | 4 | Yes |
| DS6 | 1 | Ecoli | 100 | 10 | 137 | 6 | Yes |
| DS7 | 1 | Ecoli | 100 | 10 | 137 | 7 | Yes |
| DS8 | 1 | Ecoli | 100 | 10 | 137 | 3 | Yes |

We compared several commonly used summary statistics between each synthetic data set and a matching real data set (Figure 2). These include two cell-level statistics – “library size” and “zero count per cell” (number of genes with zero recorded expression in a cell) – and three gene-level statistics – “zero count per gene” (number of cells in which a gene has zero recorded expression), “mean count” and “variance count” (mean and variance of expression of genes). As shown in Figure 2, there is strong qualitative agreement between real and synthetic (noisy) data sets in terms of each of these five statistics. As expected, the clean form of each synthetic data set has substantially different statistical properties from real data (For a more intuitive interpretation of the “total variation” metric used to compare distributions, see Supplementary Figure S2).

**Figure 2:**
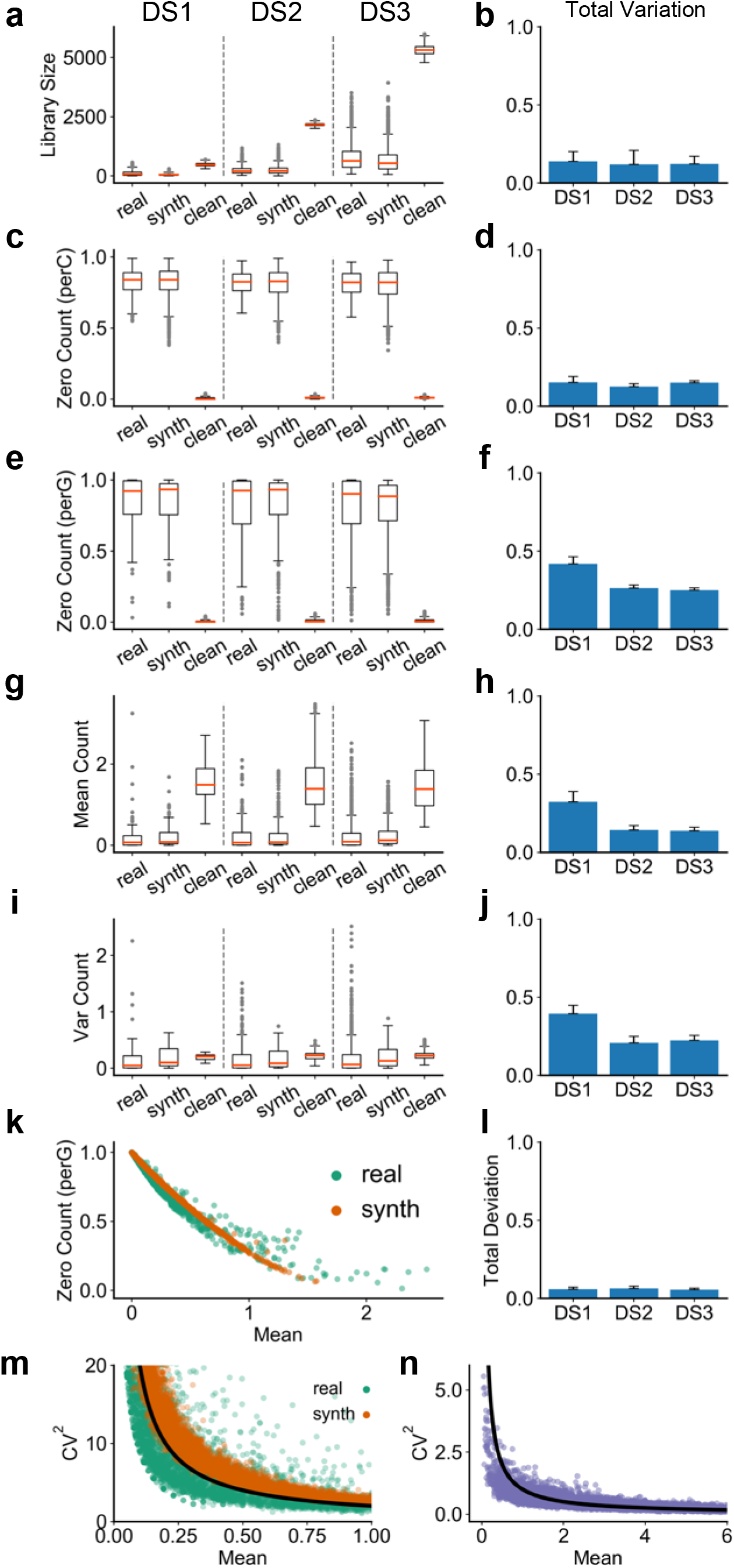
Comparisons between synthetic data generated by SERGIO and a real scRNA-seq data set. We show the distributions of per-cell quantities in (a,c), and per-gene quantities in (e, g, i), for DS1, DS2, and DS3 separated by dashed lines. These comparisons are shown between one sample from the real data set (“real”), one replicate of clean simulated data (“clean”), and its technical noise-added version (“synth”). More comprehensive comparisons – between every pair of a noisy simulated replicate and a real sample – are shown in panels to the right: the total variation (see METHODS) is calculated to compare the real and synthetic distributions and the average total variation across all such pairs is shown in panels (b, d, f, h, j). **(a,b)** Distributions and total variation of library sizes. **(c,d)** Distributions and total variation of zero counts per cell (normalized by number of genes). **(e,f)** Distributions and total variations of zero counts per gene (normalized by total number of cells). **(g,h)** Distributions and total variations of genes’ mean expression. **(i,j)** Distributions and total variations of genes’ expression variances. **(k)** Inverse relation between normalized zero count of each gene and its mean expression. Data are shown for one of the simulated replicates of DS3 and one sample from the real data containing 2500 single cells and 1200 genes selected at random. **(l)** Total deviation is calculated between two curves derived from real and synthetic points shown in (k) (see METHODS), repeated for every pair of a noisy simulated replicate and a real sample, and the average total deviation is shown. **(m)** Inverse relation between squared coefficient of variation (see METHODS) and mean expression of genes over all single-cells. Data are shown for one of the simulated replicates of DS3 and one sample from the real data containing 2500 single cells and 1200 genes selected at random. The black line shows an arbitrary function of form *y* ∼ 1/*x* which completely matches with the observed behavior in both real and synthetic data. **(n)** The inverse relation of form *y* ∼ 1/*x* is not a result of technical noise and is also observed in clean simulated data.

An empirical observation about scRNA-seq data reported in the literature is that there is an inverse relationship between the number of zeros in the recorded expression of a gene and its mean expression level across cells^49,50^. This inverse relationship is clearly seen in our (noisy) synthetic data sets and their corresponding real data sets (Figure 2k,l), and arises not only because genes with lower expression levels are more likely to result in sampled zero counts, but also because the simulator creates “dropouts” (a form of technical noise) with higher probability for such genes. Similarly, an inverse relationship between the coefficient of variation (CV) – a common measure of expression noise – and mean expression of a gene has been extensively discussed in the literature^51–53^. Figure 2m shows the existence of this relationship in a representative synthetic data set as well as in a corresponding real data set. It is not the result of adding technical noise, and is present in the clean synthetic data sets as well (Figure 2n). It arises naturally from the gene regulatory model implemented in SERGIO, in contrast to other single cell simulators that explicitly add such a relationship to their statistical sampling procedures^32^. In other words, the synthetic data sets generated by SERGIO not only exhibit realistic distributions of key summary statistics (Figure 2a-j), they also exhibit second-order relationships between pairs of variables that are characteristic of real data sets (Figure 2k-n).

### Simulated data exhibit cell heterogeneity similar to real data

Motivated by the growing use of single cell RNA-seq data to characterize cellular heterogeneity in biological samples, we next asked if the synthetic data sets from SERGIO exhibit heterogeneity similar to real ones. We first used Principal Components Analysis (PCA) to reduce each cell’s representation to 10 dimensions and then used the popular tSNE algorithm to plot cells in two dimensions. Figures 3a and 3b show such tSNE plots for a representative synthetic data set (in the DS3 setting) in their clean and noisy forms respectively. It is clear that in the absence of technical noise the nine cell types (as specified during simulation) are highly distinguishable, and that the noisy data sets smear this visual separability significantly.

**Figure 3:**
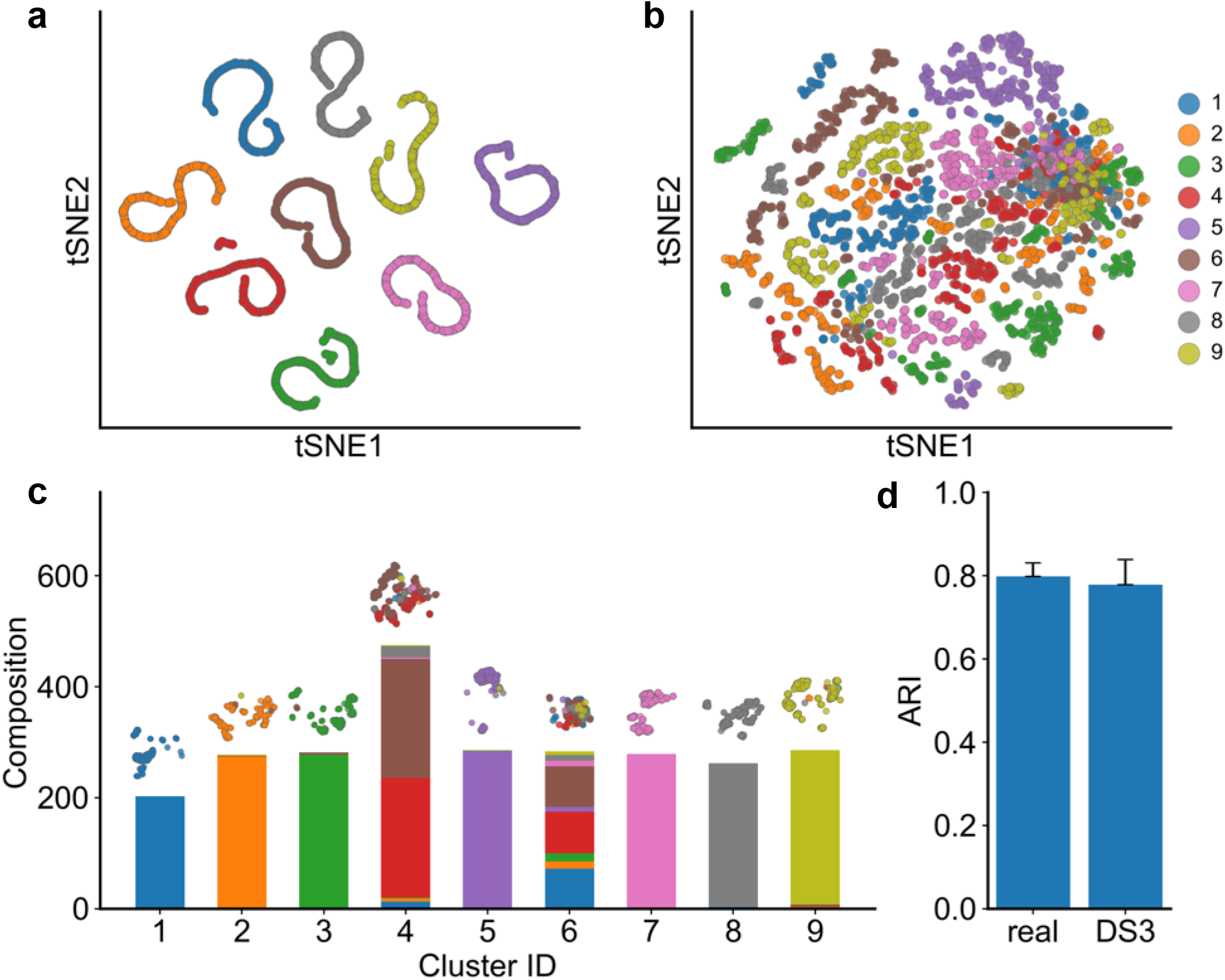
Cell heterogeneity in synthetic data generated by SERGIO. **(a)** tSNE plot of the single-cells in one simulated replicate of DS3. All cells of the same cell type are correctly clustered together. **(b)** tSNE plot of the same data set after adding technical noise. Cells are scattered such that two dimensions of tSNE representation are not sufficient for the human eye to distinguish different cell types. (The data set was not normalized for library sizes or filtered for rare genes prior to plotting.) **(c)** The noisy data set shown in (b) was clustered into nine groups using SC3 clustering method. Cell type compositions of all nine groups are shown, revealing that SC3 can correctly separate 7 of 9 cell types. Clusters 4 and 6 are less homogeneous and comprise a mixture of multiple cell types. **(d)** SC3 was applied to all 50 real samples (random subsets of the real data set), each containing 2500 cells and 1200 genes, as well as to all 15 simulated replicates of DS3. The adjusted rand index (ARI) was calculated for each clustering task, comparing the SC3 clusters to (known) true clusters defined by cell types and the average ARI is shown for each type of data (real or synthetic). ARI values obtained from simulated data are very close to those observed in real data sets.

However, cell type detection in practice does not rely only on visual separation, and specialized high-dimensional clustering algorithms are being developed for the purpose. One such algorithm is SC3^11^, which has been shown to have high accuracy for the task. It was used by Aibar et al.^9^ to cluster mouse cortex cells in the “real data set” of our study^48^ and the clusters were found to be very similar to the true cell types present in the sample (Adjusted Rand Index, ARI, of ∼0.8). If our synthetic data sets exhibit similar levels of cellular heterogeneity as the real set, then we expect SC3-reported clusters to have similar levels of concordance with “true” cell types as known to the simulator. Figure 3c shows the composition of nine clusters found by SC3 on the (noisy) synthetic data set visualized in Figure 3b, in terms of the true cell types present in each cluster. We note that seven of the nine reported clusters predominantly comprise cells of one (distinct) type, and only two of the clusters are of mixed composition, thus suggesting a high accuracy of clustering. To make this observation more formal, we computed the Adjusted Rand Index (ARI) between SC3-reported clusters and true cell types for each of the 15 replicates of the DS3 data set, noting an average ARI of 0.78. We repeated this for each of the 50 sampled subsets of the real data set corresponding to DS3 settings, and found the average ARI to be 0.80, very close to that seen in synthetic data. This exercise demonstrates that synthetic data sets generated by SERGIO exhibit realistic levels of cellular heterogeneity also illustrates the use of SERGIO to benchmark clustering methods.

### Benchmarking GRN reconstruction methods

We next illustrate how the data synthetized by SERGIO can serve to benchmark GRN reconstruction tools. In our first tests we worked with clean data sets generated by SERGIO, reasoning that these should provide an upper bound for performance on noisy realistic data sets. We evaluated the popular GRN inference algorithm called GENIE3^54^, which was originally developed for analyzing bulk RNA-seq data but has since been used successfully on single cell data as well. We applied GENIE3 on the (clean) data sets DS1 (100 genes) and DS3 (1200 genes) and evaluated the predicted TF-gene pairs based on the underlying GRNs in these data sets, using the common metrics Area Under Receiver Operating Characteristics (AUROC) and Area Under Precision-Recall Curve (AUPRC). Recall that these data sets were synthesized to include 300 cells for each of nine cell types. To assess the impact of data set size, we created smaller sets by sampling 200, 100 or 10 cells per cell type from the original simulated data (for each replicate of DS1 and DS3), and repeated the GRN reconstruction assessments for these. We also sought to assess the advantage of having single cell resolution in the data, and thus synthesized “bulk” expression data sets by averaging the expression of each gene in all cells of the same type, mimicking a situation where each cell type has been sorted separately and subjected to traditional expression profiling. (The resulting synthetic data sets included nine conditions with “bulk” expression values of each of 100 or 1200 genes, depending on the original data set.)

Figures 4a and 4b show the ROC and PRC respectively for a representative replicate of the DS3 data set, in its original setting (300 cells per type) as well as its sampled smaller versions and their respective “bulk” data set versions. A more comprehensive view, spanning all replicates of DS1 and DS3, is shown in Figure 4c-f. Several points are apparent from these figures. First, in nearly all versions of the data sets, GENIE3 performs significantly better than random, as is evident from AUROC values well above the 0.5 value expected from a random predictor. Second, we note that while performance is significantly better on larger data sets than on the smallest data set (10 cells per type), there is not a clear difference among the data sets with 100 cells per type or more. This suggests that, at least in the absence of technical noise, the benefits of greater cell count for GRN reconstruction accuracy saturate at commonly seen levels. Third, the “bulk” data sets consistently yielded lower accuracy than the single-cell data sets, regardless of the numbers of cells, confirming the value of the latter for regulatory inference. Finally, we noted that although the DS1 and DS3 data sets had similar AUROC values, the AUPRC values revealed significantly worse predictions in the larger (DS3, 1200 genes) data sets. This is expected, in part because the random baseline is lower for DS3 (random AUPRC of 0.002) than for DS1 (random AUPRC of 0.026), but also because high levels of gene co-expression confound methods such as GENIE3 more for larger data sets.

**Figure 4:**
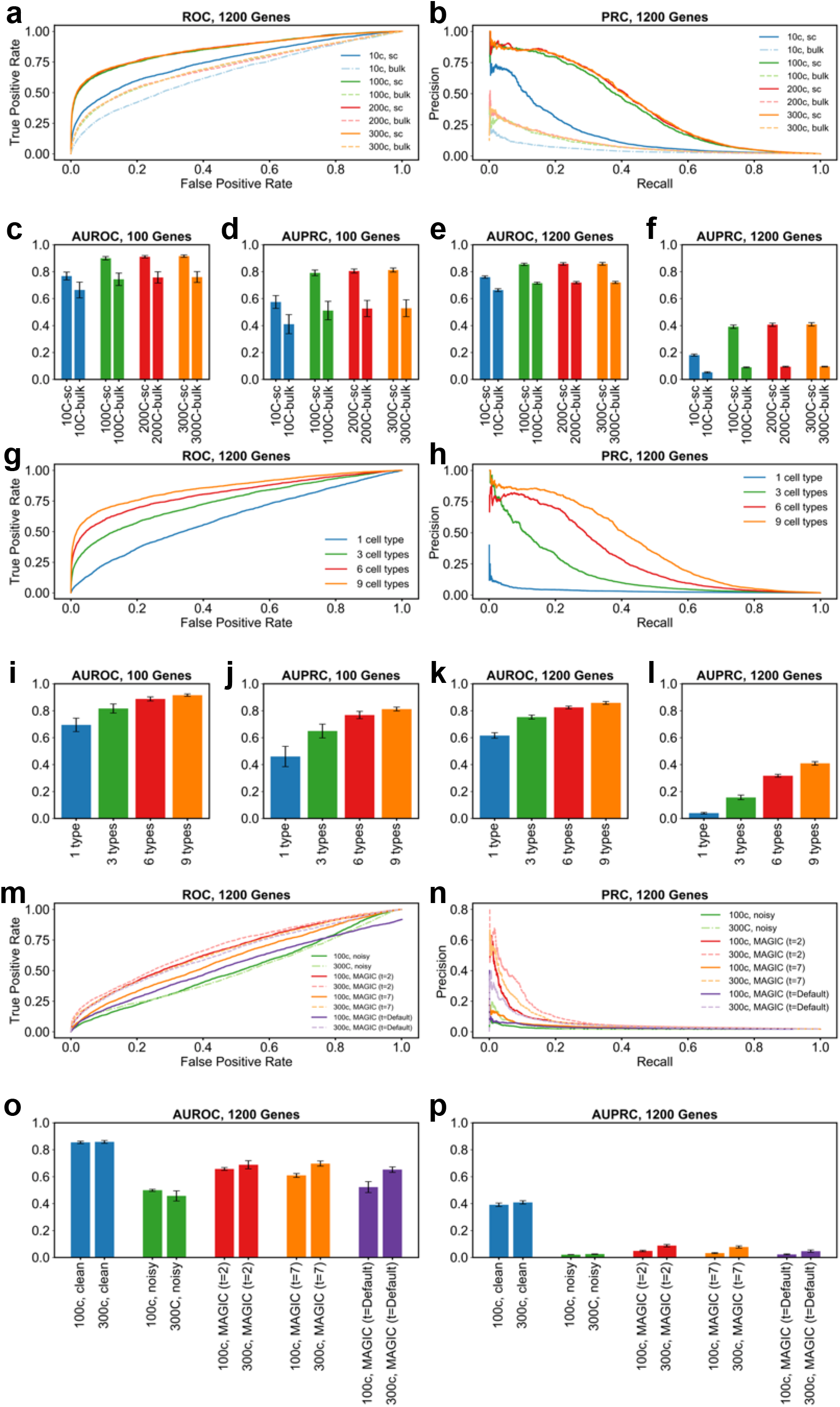
GRN inference from synthetic data generated by SERGIO. **(a,b)** Receiver Operating Characteristic (ROC) curves and Precision-Recall Curves (PRC) respectively of GRN inferred by GENIE3 on one replicate of clean synthetic data (DS3) and various subsets thereof consisting of varying number of cells per cell type. “300c” setting refers to the entire DS3 replicate, including 300 cells per cell type. The other settings (“200c”, “100c”, “10c”) refer to data sets where we sampled 200, 100, or 10 cells respectively from each cell type in DS3. For each data set, we evaluated GENIE3 on single-cell data (“sc” setting), and bulk data (“bulk” setting) which are obtained by averaging expression profiles of all cells of the same type in the corresponding single-cell data set. **(c-f)** Area under ROC curve (AUROC) and area under PR curve (AUPRC) of the GRN inferred by GENIE3, averaged over al clean replicates of DS1(c,d) or DS3 **(e,f)** and their subsets comprising varying number of cells per cell type in both single-cell and bulk settings. **(g,h)** Similar to (a,b), except that subsets of the clean synthetic data set DS3 were created to have varying numbers of cell types; every cell type retains all of its 300 simulated single-cells in the original DS3 replicate. **(i-l)** AUROC and AUPRC of GRN inference by GENIE3, averaged overall all replicates of DS1 **(i,j)** or DS3 **(k,l)**, as well as their subsets comprising varying numbers of cell types; as in (c-f), evaluations were done for the single-cell data set as well as bulk data sets derived from them. **(m,n)** ROC and PRC of the GRN inferred by GENIE3 on one noisy replicate of DS3 (“300c”) with 300 cells per type and a sub-sample of it (“100c”) containing 100 cells per type. Also shown are results of GRN inference by the same method on the same data sets, but using data imputation by MAGIC in three different settings (t=2, t=7, t=default). **(o,p)** AUROC and AUPRC of the inferred GRN by GENIE3 on all replicates of the data sets used in (m,n) as well as all the clean replicates of DS3 (“300c, clean” and “100c, clean”).

We next examined the impact of cellular heterogeneity on GRN reconstruction accuracy, using our clean synthetic data sets. For this, we sampled from each replicate of DS1 and DS3 (at their original setting of 300 cells per type) smaller data sets comprising 6, 3 or 1 cell type rather than the 9 cell types simulated. As shown via AUROC and AUPRC measures in Figures 4i-l (with representative ROC and PRC curves in Figures 4g,h), we found data sets with greater heterogeneity to consistently improve GENIE3 performance, which remained clearly above the random baseline (AUROC of 0.5 and AUPRC of 0.026 and 0.002 for DS1 and DS3 respectively) for all but the “1 cell type” setting. This is expected, since the latter setting includes gene expression variation resulting only from biological noise, and even though extrinsic noise (fluctuations in TF levels reflected in target gene levels^55^) may be exploited to infer TF-gene relationships, such correlations are diluted by the presence of intrinsic gene expression noise in the simulations (see Methods). On the other hand, in settings with 3 – 9 different cell types, the dominant form of expression variation arises from differences in the steady state profiles of the cell types, making regulatory inferences more effective.

We next examined the effect of technical noise on GRN reconstruction. For this, we compared GENIE3 performance on clean and noisy versions of each replicate of DS3 (1200 genes), in the original setting of 300 cells per type as well as a sampled version thereof with 100 cells per type. The complete results are shown in Figures 4o,p, with representative ROC and PRC curves shown in Figures 4m,n. Both performance metrics (AUROC and AUPRC) deteriorate to levels expected from random prediction when analyzing noisy synthetic data, in contrast to the very high levels seen prior to introducing technical noise. Such nearly-random performance of GENIE3 on noisy single-cell expression data has been reported in previous studies conducted based on real as well as synthetic single-cell expression data sets^43,56^. Notably, increasing the number of cells (from 100 per type to 300) does not change our conclusion.

In light of the above finding, we considered the possibility of using imputation tools specialized for single cell RNA-seq data as a means to improve the signal necessary for GRN reconstruction. We thus utilized the popular imputation tool called MAGIC^17^ to preprocess the noisy synthetic data sets prior to analyzing them with GENIE3, and compared the performance metrics to those obtained above. Results were only modestly improved from those without imputation, with AUROC values ∼ 0.65 in the 300 cell/type setting and ∼ 0.52 in the 100 cell/type setting (Figures 4m-p). Closer examination revealed that the default settings of MAGIC made the data overly structured, resulting in unrealistically large gene-gene correlations (Supplementary Figures S3 and S4), similar to previous reports^57–59^. In order to address this issue, we employed two smaller values of the ‘*t*’ parameter in MAGIC (*t* = 2 or 7), in separate runs, prior to GRN reconstruction. Both of these settings resulted in improved performance over the default setting of MAGIC, and substantially better than that seen in noisy data sets without imputation (Figures 4m-p). For instance, AUROC values for the 300 cell/type setting were at ∼0.70 (*t* = 7), squarely in the middle of those without imputation (∼0.46) and those on clean data sets (∼0.86). AUPRC values (∼0.08) were also significantly above random expectation (∼0.002), though far from the high values ∼0.4 observed on clean data sets. Although we noted above that GRN reconstruction accuracy on clean data sets did not improve when increasing the cell counts (300 versus 100 cells per type), we do notice a significant and consistent effect of cell counts in performance on imputed data (Figures 4o,p). Presumably, greater cell counts are beneficial for the imputation step, which in turn results in higher performance of GENIE3. Our overall conclusion from the above tests (Figure 4) is that a state-of-the-art GRN reconstruction method such as GENIE3^54^ can perform accurately on single cell expression data in the hypothetical scenario where technical noise is absent, but falls to near-random performance in the face of realistic levels of technical noise. The accuracy does improve above random baseline if the data are imputed with specialized tools but remains far short from the upper bar observed in clean data, making technical noise a major factor for future GRN reconstruction methods to address.

### Benchmarking differentiation trajectory inference tools

Our analysis so far involved using SERGIO to synthesize steady-state expression profiles representing different cell types. The simulator is additionally capable of synthesizing dynamic expression data on a set of genes controlled by a given regulatory network in single cells differentiating along a given trajectory (Figure 5). In this mode the simulator is provided with a differentiation graph whose nodes represent stable cell types in a differentiation program and whose edges represent differentiation from the parent cell type to child cell type. The simulator samples expression profiles from the steady state represented by the parent cell type, and then simulates a dynamical process (identical to that described above) that begins with one of these expression profiles and evolves into the steady state represented by the child cell type. It then samples expression profiles from the temporal duration when the cells are transitioning from the initial to final cell type. The entire “clean” data set is synthesized by repeating this simulation process for each edge in the differentiation graph. Technical noise is then added in a manner identical to the steady state simulation mode.

**Figure 5:**
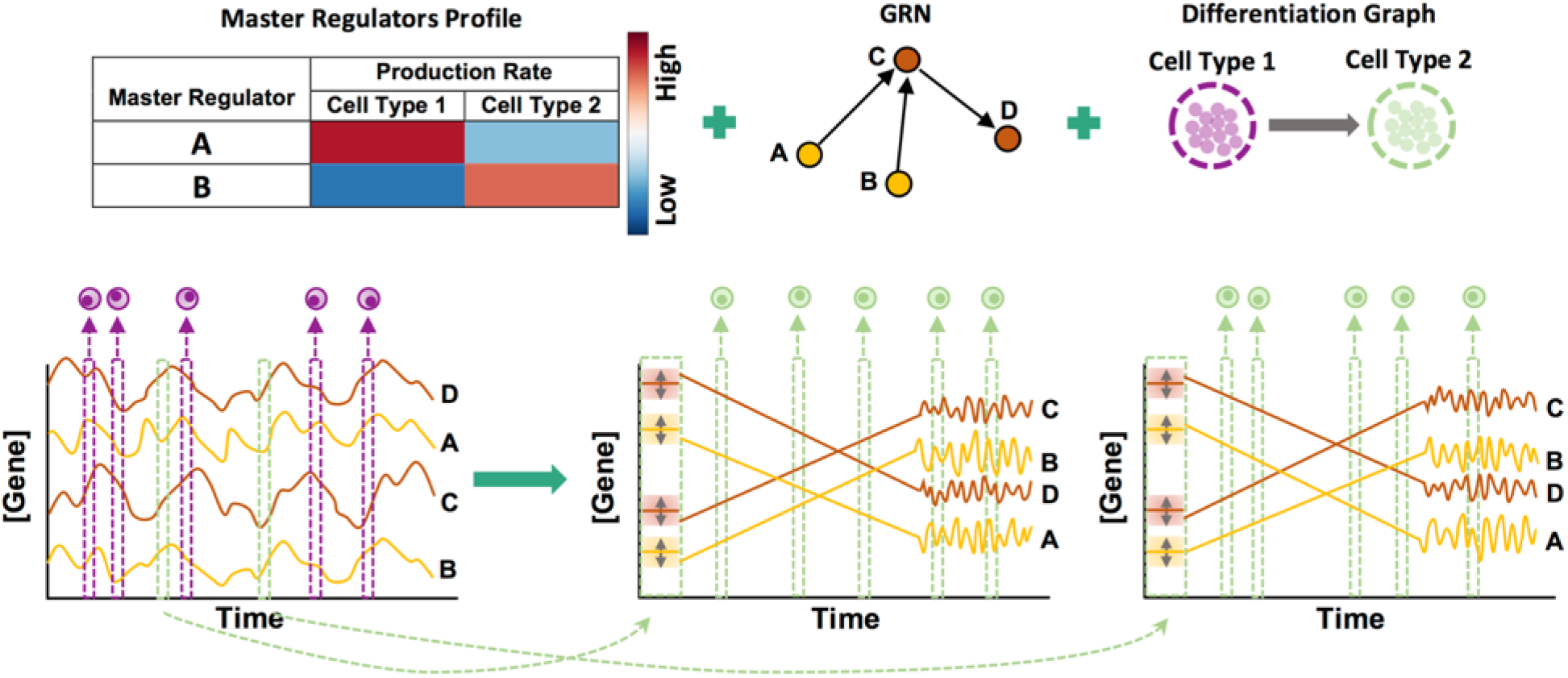
Overview of differentiation simulation pipeline. In addition to master regulators profile defining cell types and GRN, this mode requires a differentiation graph as an input (top right). Differentiation simulation is started from the origin of differentiation trajectory (cell type 1 here). Since the origin cell type is not differentiated from any other cell type, its samples (cells) are drawn from steady-state simulations (bottom left). For each child cell type (cell type 2 in this case), SERGIO initializes transcript levels to values close to their steady-state concentrations in the parent cell type (cell type 1 here). Simulations are then performed so that transcript levels reach their steady-state concentrations in the child cell type, after which simulations continue for a user-defined number of additional steps so as to collect sufficient time-course data in steady-state. SERGIO repeats simulation of the child cell type (from initialization until steady-state) for a user-defined number of times to sample enough paths between the parent and the child cell type. (Two such repeats are shown here.) Finally, single-cells belonging to the child cell type are sampled from the aggregated pool of all time-course data from the initial to the last simulation time point.

An emerging approach to describe the dynamics of differentiation programs through single-cell expression profiling involves examination of spliced as well as unspliced transcript levels in the data and inferring “RNA velocity” of each gene^60^. To allow synthesizing data sets amenable to such analysis, the differentiation simulation mode uses a variation on the underlying model described above. In particular, it invokes two chemical Langevin equations (CLE) similar to equation 1 to generate unspliced and spliced transcript levels (see Equation 8 and 9 in Methods). It reports the simulated expression values as levels of unspliced as well as spliced transcripts, whose sum may be considered the total expression of a gene.

To illustrate these features of the simulator, we generated four synthetic differentiation data sets (DS4 – DS7), each containing 100 genes controlled by the same GRN, but obeying different differentiation graphs – linear (DS4), bifurcation (DS5), trifurcation (DS6) and tree (DS7) (Figure 6, top). Figure 6a shows the two dimensional PCA plot of the clean total transcriptome (without technical noise added) for the four types of differentiation graphs. It is visually evident that these two-dimensional representations of cells based on their gene expression profiles match their corresponding graphs used in the simulations. We note that the dispersion of cells of each type (end points of each branch of a graph) as well as the width of the differentiation path from one type to another can be controlled by user-specified parameters in SERGIO (Supplementary Figure S5).

**Figure 6:**
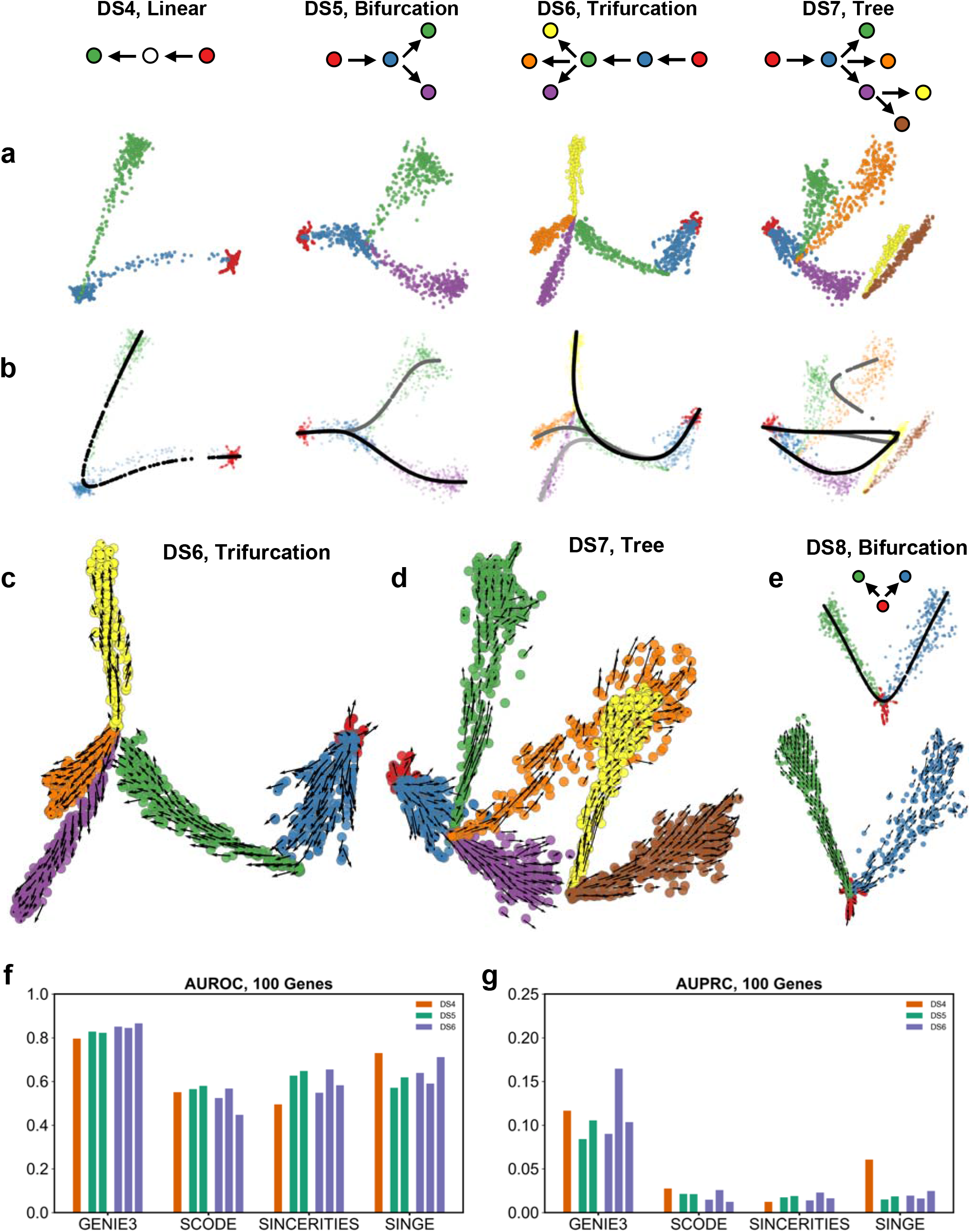
Evaluation of differentiation data sets generated by SERGIO. **(a)** PCA plots of single-cell data sets synthesized for differentiation graphs shown at top: DS4 (linear), DS5 (bifurcation), DS6 (trifurcation), and DS7 (tree). Cells of or differentiating into each cell type are shown by a distinct color. **(b)** Differentiation trajectories inferred by Slingshot on the same four data sets. Each line with a slightly different grayscale color denotes a distinct inferred path. **(c,d)** Velocity fields inferred by Velocyto on DS6 and DS7 respectively. **(e)** Differentiation trajectory inferred by Slingshot (on top) and trajectory interpreted from the velocity field inferred by Velocyto on a simple bifurcation data set (DS8) synthesize by SERGIO. **(f,g)** AUROC and AUPRC respectively of the GRN inferred by various methods on the pseudotime-ordered single cells in data sets DS4, DS5, and DS6. GRN inference was performed on each differentiation branch separately and AUROC and AUPRC is calculated and shown for each branch of DS5 and DS6.

Differentiation data sets synthesized by SERGIO can be used to benchmark trajectory inference algorithms since the underlying differentiation trajectory (graph) is known for these data. To illustrate this, we applied the Slingshot^13^ tool on the above data sets, still in their clean form without technical noise. Slingshot is a tool specifically developed for trajectory inference, with published reports of high accuracy. Consistent with these reports, we noted that Slingshot infers the correct trajectory in three of the four data sets; however, it failed to reconstruct the more complex, tree trajectory (Figure 6b) of DS7.

We then analyzed the above synthetic data sets with the Velocyto^60^ tool, which infers an “RNA velocity” field in a low dimensional representation of single cells that indicates the direction in which each cell’s expression profile appears to be changing. The velocity field also provides an intuitive visualization of differentiation trajectories. Figures 6c,d depict the inferred velocity fields for DS6 and DS7, demonstrating how Velocyto correctly captures these differentiation trajectories, including the tree of DS7 (Figure 6d) that Slingshot was unable to recover (Figure 6b, right). (Velocyto output for DS4 and DS5 may be found in Supplementary Figure S6.) Thus, we find that use of an additional layer of information – spliced versus unspliced mRNA counts – can improve trajectory inference from single cell transcriptomic data. This is not limited to data sets with complex underlying trajectories – Figure 6e shows an example data set (DS8) generated using a simple bifurcation graph for which Slingshot infers a linear trajectory while Velocyto reports a velocity field clearly indicative of the true bifurcation trajectory. It is worth noting here that the Slingshot tool may be made to utilize prior knowledge of stable cell types, and we did not provide such information, which may resolve the errors noted above. To summarize, synthetic data sets generated by SERGIO show that, at least in the absence of prior information on stable cell types, RNA velocity-based approaches may have an advantage in terms of trajectory inference on single cell data.

### Benchmarking GRN reconstruction on differentiation data

Single-cell transcriptomic profiles of differentiation processes offer unique opportunities for GRN reconstruction, where cells are ordered by “pseudotime” (a temporal partial ordering obtained by mapping them to inferred differentiation paths) and the resulting pseudotime labels are exploited to infer causal relationships between TFs and target genes. Several methods have been recently proposed that specifically channel this opportunity, including SCODE^56^, SINCERITIES^61^ and SINGE^62^. We used the dynamic data simulated by SERGIO to benchmark these specialized GRN-reconstruction algorithms, using Slingshot for pseudotime inference. We used one simulated replicate of DS4, DS5 and DS6, for which we verified above that Slingshot infers trajectories accurately. For each data set, we evaluated and compared the three above-mentioned GRN reconstruction methods on single cells associated with a single branch of the inferred differentiation trajectory (see Methods). We also used GENIE3 as a baseline method to infer TF-gene relationships without utilizing pseudotime information. Interestingly, GENIE3 clearly outperforms the three specialized algorithms in all six evaluations (Figure 6f,g). In other words, the use of temporal ordering of single cells does not help GRN reconstruction, at least in the absence of technical noise.

## Discussion

The main distinguishing quality of SERGIO is its ability to simulate single-cell expression data based on a specified GRN. Its implementation strikes a balance between a biologically realistic model of transcriptional processes and simplifying assumptions that facilitate fast simulation, capable of scaling to thousands of genes and regulatory interactions. SERGIO employs an intuitive definition of cell types as steady states of GRN dynamics^63^, and can simulate any number of user-defined cell types. It can also simulate collections of cells differentiating from one cell type to another, an important feature not available in GNW^38^ even after modifications to simulate single-cell data. Additionally, by including separate simulation of unspliced and spliced transcripts in differentiating cells, SERGIO allows assessment of tools based on the emerging approach of RNA velocity.

The unique features of SERGIO make it a powerful tool for benchmarking a wide variety of single-cell analysis tools. We have presented several examples of such benchmarking efforts, which yielded useful insights about the evaluated tools. For instance, we showed a simple example (Figure 6e) of a differentiation data set where RNA velocity-based inference outperforms alternative trajectory inference algorithms. Our assessment of a leading GRN inference tool found that it is rendered largely inaccurate (close to random performance) due to technical noise typical of contemporary data sets, even though they are capable of far greater accuracy in the absence of measurement errors. In the same context, we noted that imputation algorithms such as MAGIC^17^ can alleviate this problem to an extent, leading to modestly improved accuracy, especially if data sets have larger numbers of cells.

We also evaluated GRN inference methods designed specifically for time-ordered single-cell expression data^56,61,62^, and were surprised to find that these specialized methods are less effective than a more general-purpose method – GENIE3^54^ – even for differentiation data sets. However, the performance of these specialized tools depends on the type of differentiation trajectories, number of single-cells and other factors. For example, SINGE^62^, one of the evaluated methods, is designed to be used with an ensemble of parameter settings, and in our evaluations we used this tool with only two sets of parameters; its performance might have been significantly better if a larger ensemble of parameters were to be used.

It should be noted that the GRN benchmarking in this study considered methods based on expression only, while better accuracy can result from existing tools that use additional information such as TF-DNA binding data^9^. Future work can combine SERGIO simulations of single-cell expression with existing ideas on benchmarking GRN inference from bulk data and prior information^64^. Expression data from TF knockout experiments can also be exploited by GRN inference algorithms^65^, and knockout of master regulators (MR) can be easily simulated in SERGIO to assess such algorithms.

In conclusion, we believe that SERGIO will prove useful to a number of researchers developing tools for the rapidly developing field of single-cell transcriptomics. It will be especially useful for testing GRN reconstruction methods, which according to our assessments is the analytical task most in need of future improvements. But its usefulness will extend to future tools for other popular tasks as well, since synthetic data sets that capture real data more closely naturally provide more reliable assessments of those tools. Moreover, the “clean” simulated data sets (without technical noise) generated by SERGIO should be useful in their own right, since they also capture realistic expression variation due to biological noise and can provide upper bounds on accuracy in the idealized scenario where measurement noise has been eliminated.

## Methods

### Steady-State Simulations

We model the dynamics of the concentration of genes using systems of stochastic differential equations (SDE) that have been previously employed in GeneNetWeaver (GNW)^38,40^ and which are derived from the chemical Langevin equation (CLE)^44^. The time-course of mRNA concentration of gene *i* is modeled by:

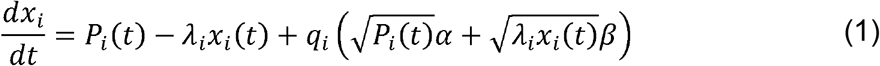

where *x_i_* is the expression of gene *i, P_i_* is its production rate, which reflects the influence of its regulators as identified by the given GRN (details below), *λ_i_* is the decay rate, and *q_i_* is the noise amplitude in the transcription of gene *i. α* and *β* are two independent Gaussian white noise processes. In order to obtain the mRNA concentrations as a function of time, we integrate the above stochastic differential equation for all genes:

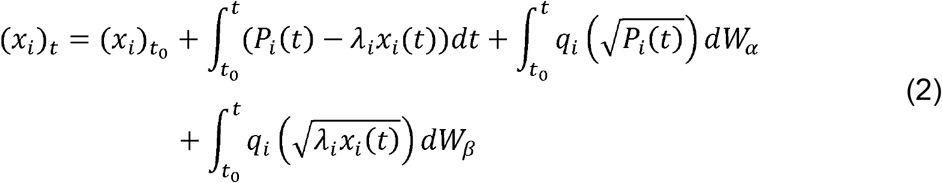

where *W_α_* and *W_β_* are two independent stochastic Wiener processes. We integrate this equation in pre-defined time steps of duration *Δt*, according to Euler–Maruyama method^66^ using the Itô scheme:

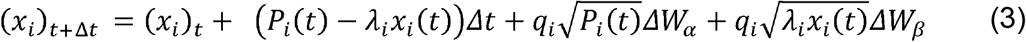

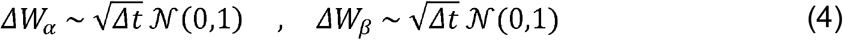

Each iteration yields the mRNA concentrations of all genes at time step *t* + Δ*t* using each gene’s own concentration and all of its regulators’ concentrations at time step *t*.

We model each gene’s production rate, *P_i_*, as the sum of contributions from each of its regulators (as prescribed by the GRN):

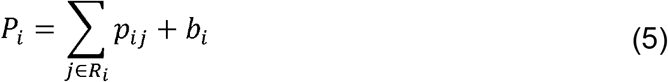

where *R_i_* is the set of all regulators of gene *i, b_i_* is the basal production rate of gene *i*, and *p_ij_* is the regulatory effect of gene (TF) *j* on gene *i*. The latter is modeled as a nonlinear saturating Hill function of the mRNA concentration of the TF^46^:

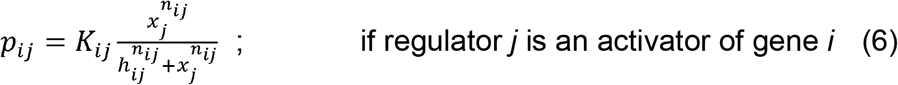

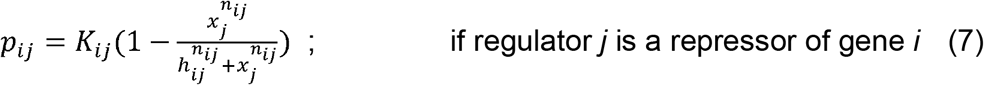

where *K_ij_* denotes the maximum contribution of regulator *j* to target gene *i, n_ij_* is the Hill coefficient that introduces non-linearity to the model and *h_ij_* is the regulator concentration that produces half-maximal regulatory effect (half-response). If gene *i* is a user-designated “master regulator” (MR), i.e., no gene regulates it, then its production rate *P_i_* is entirely determined by basal production rate *b_i_* which is a user-defined parameter. For simplicity, we set *b_i_* = 0 for genes other than master regulators. *K_ij_* and *n_ij_* are user-defined parameters, and the type of each interaction (activation or repression) is also user-specified. The *h_ij_* parameter is set to be the average of the regulators’ expression among the cell types to be simulated. The parameters *α* and *β* in equation 1 characterize the intrinsic noise associated with the production and decay processes of the mRNA transcript of gene *i*. Moreover, the intrinsic noise in the transcription of regulators propagates along the GRN and thus influences the production rate *P_i_* to become an extrinsic noise source in the transcription of gene *i*. We support three forms of noise:

1. Dual Production Decay (“dpd”): the form of stochastic noise that is shown in equation 1.
2. Single Production (“sp”): including only the noise term associated with the production process (equivalently, *β* = 0 set).
3. Single Decay (“sd”): including only the noise term associated with the decay process (equivalently, set *α* = 0)

We note that the current version of Sergio is not capable of simulating GRNs containing auto-regulatory edges or cycles.

### Sampling Single Cells

We use the above system of equations to simulate the time-course of each gene’s expression in a cell, starting with a given initial value, and record expression values of all genes at randomly selected time points after the simulation has reached steady state. Invoking the ergodic assumption^47^, we treat the expression profiles at these time points to represent single-cell profiles. In order to speed up the simulation, we estimate the steady-state concentrations of all genes given the input parameters (see Supplementary Notes 1) and initialize the time-course simulation with those values. Also, we ensure that a sufficient number of time steps, which is controlled by a user-defined parameter, are simulated in the steady state prior to sampling cells.

### Cell Types

The above simulation is performed for each “cell type” separately. We define a cell type (or cell state) by the average concentration of master regulators. A cell type differs from another cell type by the average concentration of one or more of the master regulators among the population of cells belong to each cell type. This can be controlled by the basal production rate *b* for master regulators (see Supplementary Notes 1). Sergio takes as input the basal production rate of all master regulators in each of the cell types to be simulated.

### Simulation of differentiation trajectories

In addition to simulating one or more “cell types” in steady state, Sergio may be used to simulate cells on the differentiation trajectory from one cell type to another, i.e., between two steady states. More generally, given a “differentiation graph” where nodes represent cell types and directed edges indicate differentiation from one cell type to the other, Sergio can simulate expression profiles of cells spanning different stages of differentiation specified by the graph. Such cells are either in one of the steady states represented by nodes or have departed away from the steady-state of their “parent” cell type of an edge and are migrating toward the steady-state of the corresponding “child” cell type. The differentiation is presumed to commence when one or more master regulators change their expression from that in the steady state of the parent cell type, e.g., due to a signaling event^67^ or due to a noise-driven switch^68^. Thus, given a differentiation graph and average expression levels of master regulators for each cell type (nodes), we simulate each differentiation trajectory (edge) as follows: 1) Cells representing the parent cell type are sampled from the corresponding steady state. 2) Production rates (*P_i_*) of master regulators are changed from those specified for the parent cell type to those of the child cell type, and time-course simulations are performed following equations 3–4 as explained above. As these simulations proceed, all genes ultimately converge to their steady-state concentrations in the child cell type. 3) Cells (expression profiles) are sampled at random from the entire simulation, including cells in the parent and child cell types (steady states) as well as cells on the differentiation trajectory (transient states). Multiple such time-course simulations are performed and the sampled cells are randomly chosen from the entire collection of such simulations. Also, after each simulation reaches the steady-state of the child cell type, it may be continued for a user-defined number of additional steps. This controls the ratio of the cells in the steady states of the differentiation graph to the number of cells in differentiating (transient) states.

Simulations of differentiation trajectories in Sergio generate not only the total mRNA concentration of each gene (in a time-course), but the changing levels of spliced and unspliced mRNA transcripts separately. To this end, we express the rate of change in the concentration of unspliced and spliced RNA using ordinary differential equations (ODEs), following prior work^60,69^. Furthermore, we introduce noise terms to these ODEs in a manner similar to steady-state simulations (equation 1). Thus, the time-course of the spliced (*s*) and unspliced (*u*) transcript level of gene *i* is modeled as:

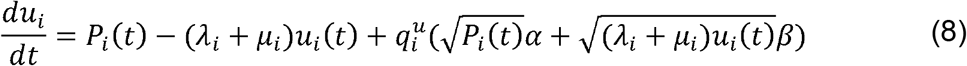

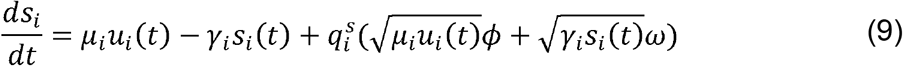

where *P_i_*(*t*) is the production rate of pre-mRNA (unspliced transcript) that includes regulatory interactions, *λ_i_* and *μ_i_* are the degradation and splicing rate respectively of pre-mRNA and 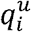 is the noise amplitude associated with the transcription of pre-mRNA. For simplicity, we assume the degradation rate *λ* of pre-mRNA is zero and all of its decay is due to splicing (user-defined parameter *μ_i_*). Also, *γ_i_* is the degradation rate of spliced mRNA and 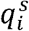 is the noise amplitude associated with the transcription of spliced mRNA. *α, β, ϕ* and *ω* are independent Gaussian white noise processes. All the three form of stochastic noise (“dpd”, “sp”, “sd”) described for steady-state simulations are also supported in dynamics simulation. Moreover, production-rate *P_i_* is modeled as in steady-state simulations (equations 5–7 above). Both of the SDEs in equations 8–9 are integrated according to Euler–Maruyama scheme to obtain time-courses of unspliced and spliced mRNA concentrations.

### Technical Noise

Sergio adopts methods similar to Splatter^32^ for adding technical noise to the simulated single-cell expression data. One module introduces the phenomenon of “outlier genes”, which refers to the empirical observation that a small set of genes appear to have unusually high expression measurements across cells in typical scRNA-seq data sets. A second module incorporates the noted phenomenon of different cells having different total counts (library size), that follows a log-normal distribution. A third module introduces “dropouts”, which refers to the observation that a high percentage of genes are recorded at zero expression in any given cell, indicating an experimental failure to record their expression rather than true non-expression. These three modules may be invoked optionally and in any combination and order specified by the user. We elaborate on each of these modules below. We focus on details pertinent to simulation of steady state data; corresponding details for differentiation trajectory data are provided in Supplementary Notes 3. Each of the modules outlined below adds a single type of technical noise to the data set provided to it.

#### Outlier genes

Each gene is designated as an outlier with a user-defined probability. If so, its expression (in every cell) is multiplied by a factor sampled from a log-normal distribution, otherwise the expression is left unchanged:

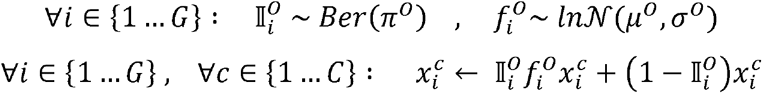

where *G* and *C* denote the total number of simulated genes and cells respectively, and 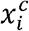 denotes the simulated expression of gene *i* in cell *c*. 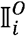 is a binary variable indicating if gene *i* is an outlier, and is sampled from a Bernoulli distribution with parameter *π^o^*.

Also, *μ^o^* and *σ^o^* are user-defined mean and standard deviation of the lognormal distribution from which the outlier scaling factor 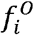 is sampled.

#### Library size

For every cell (library) a library size parameter is sampled from a lognormal distribution, and expression values of all genes in the cell are scaled by a constant factor such that the resulting total cell depth matches the sampled library size:

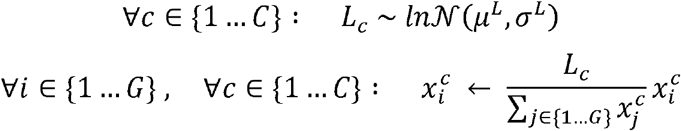

where *μ^L^* and *σ_L_* are the user-defined mean and standard deviation of the lognormal distribution from which the desired library size *L_c_* of cell *c* is sampled.

#### Dropout

To introduce dropouts to the simulated data, we first assign a probability to the expression of each gene in each of the simulated cells being a dropout. This probability is modeled as a logistic function of the expression of the gene in that cell, so that a high expression value is less likely to be zeroed out. This probability is then used as the parameter of a Bernoulli distribution from which a binary variable is sampled to indicate whether the gene is a dropout in the cell:

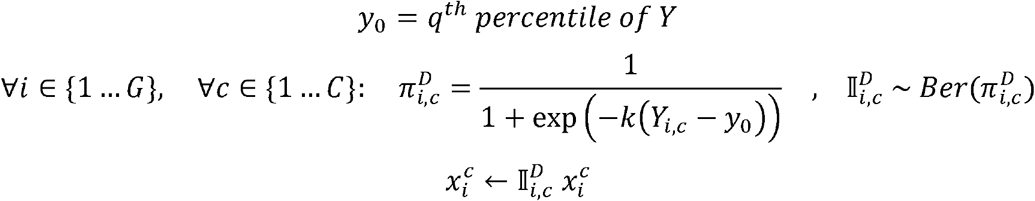

where *Y* is the expression matrix in logarithmic scale:

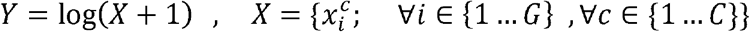

Also, *k* and *q* are two user-defined parameters that determine the logistic probability *π^D^*.

#### Conversion to UMI counts

We generate UMI counts (*UC*) by sampling from a Poisson distribution whose mean is the simulated expression level of the gene in the cell:

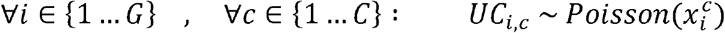

### Data Set Generation

We now describe how we set simulation parameters to generate the data sets analyzed in this study.

We sampled four gene regulatory networks (GRNs) from the known regulatory networks in S. cerevisiae and E. coli using GNW^38^ using the “random seed” argument to select genes and the “random among top 20%” setting for neighbor selection. Two of the networks consist of 100 genes and were separately sampled from Ecoli, a third network containing 1200 genes was sampled from E. coli, and the fourth network comprising 400 genes was sampled from S. cerevisiae. We also used GNW to designate each TF-gene edge as either an activating or a repressive interaction. Auto-regulatory edges were removed from the sampled networks and cycles were broken at a randomly selected edge, since Sergio does not support these two graph properties. These four networks were used to simulate 8 data sets, each with 9 cell types and 300 cells per cell type (Table 1). Fifteen “replicates” of each data set were created that had identical simulation parameters and differed due to the stochastic noise and random sampling. For all data sets, interaction strengths *K_ij_* (equations 6–7) were uniformly sampled from the range 1 to 5. Each cell type to be simulated was specified by the expression state (high or low) of each master regulator (MR); the basal production rate (*b_i_* in equation 5) of each MR was sampled from a pre-defined range that depends on the expression state and varies among different data sets (see Supplementary Table S3). We used a hill coefficient of 2 for all interactions in all data sets. We used the same noise amplitude parameter *q* = 1 and the same decay parameter *λ* = 0.8 for all genes in all steady-state data sets. In dynamics simulations, we used an unspliced noise parameter *q^u^* = 0.3 and a spliced noise parameter *q^s^* = 0.07 for all genes. Also, we used an unspliced transcript decay rate of *μ* = 0.8 and a spliced transcript decay rate of *γ* = 0.2 that maintains a ratio of spliced to unspliced expression of a gene at ∼4 (see Supplementary Notes 2). We used “dpd” setting of intrinsic noise and an integration time step of 0.01 for both steady-state and dynamics simulations.

We compared the simulated expression matrix (genes × cells) to a single-cell RNA-seq data set from the mouse cerebral cortex^48^, referred to as the “real data set”, to demonstrate that the simulated and real data sets have similar statistical properties. The real data set includes 3005 cells from nine cell types and our simulations therefore used nine cell types. However, the real data set has variable numbers of cells per cell type while we sought to keep this number fixed, or at least comparable, across cell types for ease of downstream interpretations. Hence we simulated 300 cells for each cell type (total of 2700 cells) and sampled the real data set by drawing cells of each type at random: for cell types with less than 300 cells, we retained all the cells, while for the other cell types we randomly sampled 300 cells such that a total of 2500 single cells were sampled. Our simulations generated expression values for 100, 400 or 1200 genes depending on the data set, hence we randomly sampled from the real data set the same number of genes as present in the synthetic data.

To add technical noise we used the above-mentioned modules for outlier genes, library size effect and dropouts in that order, and finally converted the expression levels to UMI counts. For each data set, we manually tuned the input parameters (see Supplementary Table S1) to each of the technical noise modules to obtain a match between the synthetic and real data. Furthermore, we filtered cells from the synthetic data that have total UMI count (sum over all genes) less than 5. In this study, we only added technical noise to the steady-state synthetic data sets and the dynamics simulations only utilized “clean data sets” without technical noise.

### Settings of single-cell analysis tools

In this study we applied several tools to the real or synthetic data sets to mimic real-world analysis of such data and to benchmark these tools. We did not normalize the data sets prior to using these tools unless otherwise specified. We note below the specific settings used for each of the tools we tested:

#### SC3^11^

We did not run SC3 to infer the number of cell types, instead we treated the number of cell types as a known quantity and required SC3 to cluster data sets into 9 cell types.

#### GENIE3^54^

We provided the identities of true regulators to the GENIE3 tool except when analyzing the differentiation data sets where we used all the genes as potential regulators. Also, for differentiation analysis GENIE3 was run on the exact same expression matrices as used for the other GRN inference tools in this study.

#### MAGIC^17^

Prior to running MAGIC, we filtered the synthetic data for rare genes (those expressed in less than 5 cells), and performed library size normalization as well as a square root transformation. We used MAGIC with the parameter *t* = 2, *t* = 7, and default setting where *t* is inferred from data.

#### SLINGSHOT^13^

We used the first two PCs as a low-dimensional representation of single-cells, and provided these as input to SLINGSHOT, along with the cell type labels. We did not provide any further prior information about origin and end cell types of trajectories.

#### VELOCYTO^60^

We performed all of the filtering and normalizations for spliced and unspliced counts that are recommended by developers of the software. We also performed K-nearest neighbor imputation on 20-dimensional PCA representations of single cells with K = 5. The “constant velocity” model was employed for inferring velocity fields, and square root transformation was used for estimating transition probabilities from PCA representation of single-cells.

#### SCODE^56^

For each differentiation trajectory, we used SCODE with *D* = 2, 4, 6, 8, and 10 to infer regulatory relationships and report the results that produce the highest

#### AUROC

Also, for each differentiation trajectory and setting of *D* parameter, we ran SCODE for 100 iterations and averaged results over 5 trials as recommended by the tool’s developers. All genes were considered as potential regulators. The inferred sign of interactions (activating or repressing) was ignored in evaluation of the tool’s performance: we sorted all gene-gene interactions by the absolute value of their inferred scores and assessed this ranked list for accuracy. Thus, the reported performance values are an overestimate of GRN inference accuracy in our setting.

#### SINCERITIES^61^

For each differentiation trajectory, we used the tool with parameters specifying Kolmogorov-Smirnov distance, Ridge regularization, and no auto-regulatory edge setting for unsigned GRN inference. As for SCODE, performance evaluation ignored signs in the true GRN.

#### SINGE^62^

For each differentiation trajectory, we used the tool with two different settings of hyper-parameters, with two replicates for each setting (for more details, see Supplementary Table S2). The best results from these four runs were reported.

#### Data sets for evaluating GRN inference on differentiation data

For each differentiation simulation data set, we generated sub-matrices that represent cells belonging to a single lineage. Therefore, we obtained 1, 2, and 3 sub-matrices for DS4, 5, and 6 respectively. Assignment of cells to different lineages was performed according to the pseudotime inferred by Slingshot, and the assigned sets of cells need not be mutually exclusive (i.e., some single cells might belong to more than one lineage). GRN inference was performed for each of the lineages separately.

### Technical Definitions

#### Total Variation (TV)

It is a measure for the distance between two probability distributions. For two probability distributions *P* and *Q* over a finite countable set *X*, total variation is defined as:

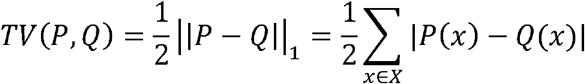

where ||.||_1_ denotes the L1 norm. Note that total variation varies in range [0 1].

#### Total Deviation (TD)

It is a measure for evaluating the difference between two functions of the same variable. For two bounded continuous function *F* and *G*, the normalized total deviation in range [a b] is defined as:

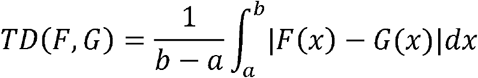

Note that if *F* and *G* are lower-bounded by zero and upper-bounded by *h*, the normalized total deviation *TD*(*F, G*) is also bounded similarly.

#### Coefficient of Variation (CV)

Characterizes the dispersion of data around its mean and is defined as the ratio of the standard deviation to the mean:

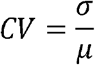

### Code Availability

SERGIO is available as a python package on GitHub: (https://github.com/PayamDiba/SERGIO).

## References

1. Papalexi, E. & Satija, R. Single-cell RNA sequencing to explore immune cell heterogeneity. Nat. Rev. Immunol. 18, 35–45 (2018).

2. Kelsey, G., Stegle, O. & Reik, W. Single-cell epigenomics: Recording the past and predicting the future. Science (80-.). 358, 69–75 (2017).

3. Park, J. et al. Single-cell transcriptomics of the mouse kidney reveals potential cellular targets of kidney disease. Science (80-.). 360, 758–763 (2018).

4. Hedlund, E. & Deng, Q. Single-cell RNA sequencing: Technical advancements and biological applications. Mol. Aspects Med. 59, 36–46 (2018).

5. Butler, A., Hoffman, P., Smibert, P., Papalexi, E. & Satija, R. Integrating single-cell transcriptomic data across different conditions, technologies, and species. Nat. Biotechnol. 36, 411–420 (2018).

6. Buettner, F. et al. Computational analysis of cell-to-cell heterogeneity in single-cell RNA-sequencing data reveals hidden subpopulations of cells. Nat. Biotechnol. 33, 155–160 (2015).

7. Stegle, O., Teichmann, S. A. & Marioni, J. C. Computational and analytical challenges in single-cell transcriptomics. Nat. Rev. Genet. 16, 133–145 (2015).

8. Wolf, F. A., Angerer, P. & Theis, F. J. SCANPY: Large-scale singlecell gene expression data analysis. Genome Biol. 19, (2018).

9. Aibar, S. et al. SCENIC: Single-cell regulatory network inference and clustering. Nat. Methods 14, 1083–1086 (2017).

10. Satija, R., Farrell, J. A., Gennert, D., Schier, A. F. & Regev, A. Spatial reconstruction of single-cell gene expression data. Nat. Biotechnol. 33, 495–502 (2015).

11. Kiselev, V. Y. et al. SC3: Consensus clustering of single-cell RNA-seq data. Nat. Methods 14, 483–486 (2017).

12. Herring, C. A. et al. Unsupervised Trajectory Analysis of Single-Cell RNA-Seq and Imaging Data Reveals Alternative Tuft Cell Origins in the Gut. Cell Syst. 6, 37–51.e9 (2018).

13. Street, K. et al. Slingshot: Cell lineage and pseudotime inference for single-cell transcriptomics. BMC Genomics 19, (2018).

14. Chan, T. E., Stumpf, M. P. H. & Babtie, A. C. Gene Regulatory Network Inference from Single-Cell Data Using Multivariate Information Measures. Cell Syst. 5, 251–267.e3 (2017).

15. Mohammadi, S., Ravindra, V., Gleich, D. F. & Grama, A. A geometric approach to characterize the functional identity of single cells. Nat. Commun. (2018). doi:10.1038/s41467-018-03933-2

16. Li, W. V. & Li, J. J. An accurate and robust imputation method scImpute for single-cell RNA-seq data. Nat. Commun. 9, (2018).

17. van Dijk, D. et al. Recovering Gene Interactions from Single-Cell Data Using Data Diffusion. Cell 174, 716–729.e27 (2018).

18. Eraslan, G., Simon, L. M., Mircea, M., Mueller, N. S. & Theis, F. J. Single-cell RNA-seq denoising using a deep count autoencoder. Nat. Commun. 10, (2019).

19. Schaum, N. et al. Single-cell transcriptomics of 20 mouse organs creates a Tabula Muris. Nature 562, 367–372 (2018).

20. Hou, R., Denisenko, E. & Forrest, A. R. R. scMatch: a single-cell gene expression profile annotation tool using reference datasets. Bioinformatics (2019). doi:10.1093/bioinformatics/btz292

21. Van den Berge, K. et al. Observation weights unlock bulk RNA-seq tools for zero inflation and single-cell applications. Genome Biol. 19, (2018).

22. Chen, C. et al. scRMD: Imputation for single cell RNA-seq data via robust matrix decomposition. bioRxiv 459404 (2018). doi:10.1101/459404

23. Gong, W. et al. A novel algorithm for the collective integration of single cell RNA-seq during embryogenesis. bioRxiv 543314 (2019). doi:10.1101/543314

24. Korthauer, K. D. et al. A statistical approach for identifying differential distributions in single-cell RNA-seq experiments. Genome Biol. 17, (2016).

25. Risso, D., Perraudeau, F., Gribkova, S., Dudoit, S. & Vert, J. P. A general and flexible method for signal extraction from single-cell RNA-seq data. Nat. Commun. 9, (2018).

26. Campbell, K. R. & Yau, C. Uncovering pseudotemporal trajectories with covariates from single cell and bulk expression data. Nat. Commun. 9, (2018).

27. Zhang, X., Xu, C. & Yosef, N. Simulating multiple faceted variability in single cell RNA sequencing. Nat. Commun. 10, (2019).

28. Marouf, M. et al. Realistic in silico generation and augmentation of single cell RNA-seq data using Generative Adversarial Neural Networks. bioRxiv (2018). doi:10.1101/390153

29. Li, W. V. & Li, J. J. A statistical simulator scDesign for rational scRNA-seq experimental design. Bioinformatics 35, i41–i50 (2019).

30. Vieth, B., Ziegenhain, C., Parekh, S., Enard, W. & Hellmann, I. powsimR: power analysis for bulk and single cell RNA-seq experiments. Bioinformatics 33, 3486–3488 (2017).

31. Papadopoulos, N., Gonzalo, P. R. & Söding, J. PROSSTT: probabilistic simulation of single-cell RNA-seq data for complex differentiation processes. Bioinformatics (2019). doi:10.1093/bioinformatics/btz078

32. Zappia, L., Phipson, B. & Oshlack, A. Splatter: Simulation of singlecell RNA sequencing data. Genome Biol. 18, (2017).

33. Intosalmi, J., Mannerstrom, H., Hiltunen, S. & Lahdesmaki, H. SCHiRM: Single Cell Hierarchical Regression Model to detect dependencies in read count data. bioRxiv 335695 (2018). doi:10.1101/335695

34. El Samad, H., Khammash, M., Petzold, L. & Gillespie, D. Stochastic modelling of gene regulatory networks. Int. J. Robust Nonlinear Control 15, 691–711 (2005).

35. Wilkinson, D. J. Stochastic modelling for quantitative description of heterogeneous biological systems. Nature Reviews Genetics 10, 122–133 (2009).

36. Kepler, T. B. & Elston, T. C. Stochasticity in transcriptional regulation: Origins, consequences, and mathematical representations. Biophys. J. 81, 3116–3136 (2001).

37. Karlebach, G. & Shamir, R. Modelling and analysis of gene regulatory networks. Nat. Rev. Mol. Cell Biol. 9, 770–780 (2008).

38. Schaffter, T., Marbach, D. & Floreano, D. GeneNetWeaver: In silico benchmark generation and performance profiling of network inference methods. Bioinformatics 27, 2263–2270 (2011).

39. Bellot, P., Olsen, C., Salembier, P., Oliveras-Vergés, A. & Meyer, P. E. NetBenchmark: A bioconductor package for reproducible benchmarks of gene regulatory network inference. BMC Bioinformatics 16, (2015).

40. Marbach, D. et al. Revealing strengths and weaknesses of methods for gene network inference. Proc. Natl. Acad. Sci. 107, 6286–6291 (2010).

41. Siegenthaler, C. & Gunawan, R. Assessment of network inference methods: How to cope with an underdetermined problem. PLoS One 9, (2014).

42. Saelens, W., Cannoodt, R. & Saeys, Y. A comprehensive evaluation of module detection methods for gene expression data. Nat. Commun. 9, (2018).

43. Chen, S. & Mar, J. C. Evaluating methods of inferring gene regulatory networks highlights their lack of performance for single cell gene expression data. BMC Bioinformatics 19, (2018).

44. Gillespie, D. T. Chemical Langevin equation. J. Chem. Phys. 113, 297–306 (2000).

45. Kolodziejczyk, A. A., Kim, J. K., Svensson, V., Marioni, J. C. & Teichmann, S. A. The Technology and Biology of Single-Cell RNA Sequencing. Mol. Cell 58, 610–620 (2015).

46. Chu, D., Zabet, N. R. & Mitavskiy, B. Models of transcription factor binding: Sensitivity of activation functions to model assumptions. J. Theor. Biol. 257, 419–429 (2009).

47. Prill, R. J., Vogel, R., Cecchi, G. A., Altan-Bonnet, G. & Stolovitzky, G. Noise-driven causal inference in biomolecular networks. PLoS One 10, (2015).

48. Zeisel, A. et al. Cell types in the mouse cortex and hippocampus revealed by single-cell RNA-seq. Science (80-.). 347, 1138–1142 (2015).

49. Pierson, E. & Yau, C. ZIFA: Dimensionality reduction for zero-inflated single-cell gene expression analysis. Genome Biol. 16, (2015).

50. Andrews, T. S. & Hemberg, M. M3Drop: dropout-based feature selection for scRNASeq. Bioinformatics (2018). doi:10.1093/bioinformatics/bty1044

51. McCarthy, D. J., Chen, Y. & Smyth, G. K. Differential expression analysis of multifactor RNA-Seq experiments with respect to biological variation. Nucleic Acids Res. 40, 4288–4297 (2012).

52. Franz, K., Singh, A. & Weinberger, L. S. Lentiviral vectors to study stochastic noise in gene expression. Methods Enzymol. 497, 603–622 (2011).

53. Dar, R. D. et al. Transcriptional bursting explains the noise-versus-mean relationship in mRNA and protein levels. PLoS One 11, (2016).

54. Huynh-Thu, V. A., Irrthum, A., Wehenkel, L. & Geurts, P. Inferring regulatory networks from expression data using tree-based methods. PLoS One 5, (2010).

55. Swain, P. S., Elowitz, M. B. & Siggia, E. D. Intrinsic and extrinsic contributions to stochasticity in gene expression. Proc. Natl. Acad. Sci. 99, 12795–12800 (2002).

56. Matsumoto, H. et al. SCODE: An efficient regulatory network inference algorithm from single-cell RNA-Seq during differentiation. Bioinformatics 33, 2314–2321 (2017).

57. Zhang, L. & Zhang, S. Comparison of computational methods for imputing single-cell RNA-sequencing data. IEEE/ACM Transactions on Computational Biology and Bioinformatics (2018). doi:10.1109/TCBB.2018.2848633

58. Peng, T., Zhu, Q., Yin, P. & Tan, K. SCRABBLE: single-cell RNA-seq imputation constrained by bulk RNA-seq data. Genome Biol. 20, 88 (2019).

59. Iacono, G., Massoni-Badosa, R. & Heyn, H. Single-cell transcriptomics unveils gene regulatory network plasticity. Genome Biol. 20, (2019).

60. La Manno, G. et al. RNA velocity of single cells. Nature 560, 494–498 (2018).

61. Papili Gao, N., Ud-Dean, S. M. M., Gandrillon, O. & Gunawan, R. SINCERITIES: Inferring gene regulatory networks from time-stamped single cell transcriptional expression profiles. Bioinformatics 34, 258–266 (2018).

62. Deshpande, A., Chu, L.-F., Stewart, R. & Gitter, A. Network Inference with Granger Causality Ensembles on Single-Cell Transcriptomic Data. bioRxiv 534834 (2019). doi:10.1101/534834

63. Huang, S., Eichler, G., Bar-Yam, Y. & Ingber, D. E. Cell fates as highdimensional attractor states of a complex gene regulatory network. Phys. Rev. Lett. 94, (2005).

64. Siahpirani, A. F. & Roy, S. A prior-based integrative framework for functional transcriptional regulatory network inference. Nucleic Acids Res. 45, (2017).

65. Bonneau, R. et al. The inferelator: An algorithn for learning parsimonious regulatory networks from systems-biology data sets de novo. Genome Biol. 7, (2006).

66. Schaffter, T. Numerical integration of SDEs: a short tutorial. Swiss Fed. Inst. Technol. Lausanne (… 1–8 (2010).

67. Basson, M. A. Signaling in cell differentiation and morphogenesis. Cold Spring Harbor perspectives in biology 4, (2012).

68. Balázsi, G., Van Oudenaarden, A. & Collins, J. J. Cellular decision making and biological noise: From microbes to mammals. Cell 144, 910–925 (2011).

69. Svensson, V. & Pachter, L. RNA Velocity: Molecular Kinetics from Single-Cell RNA-Seq. Mol. Cell 72, 7–9 (2018).

