## Supplementary Information for "A single-cell expression simulator guided by gene regulatory networks"

### Supplementary Figures

#### Supplementary Figure S1

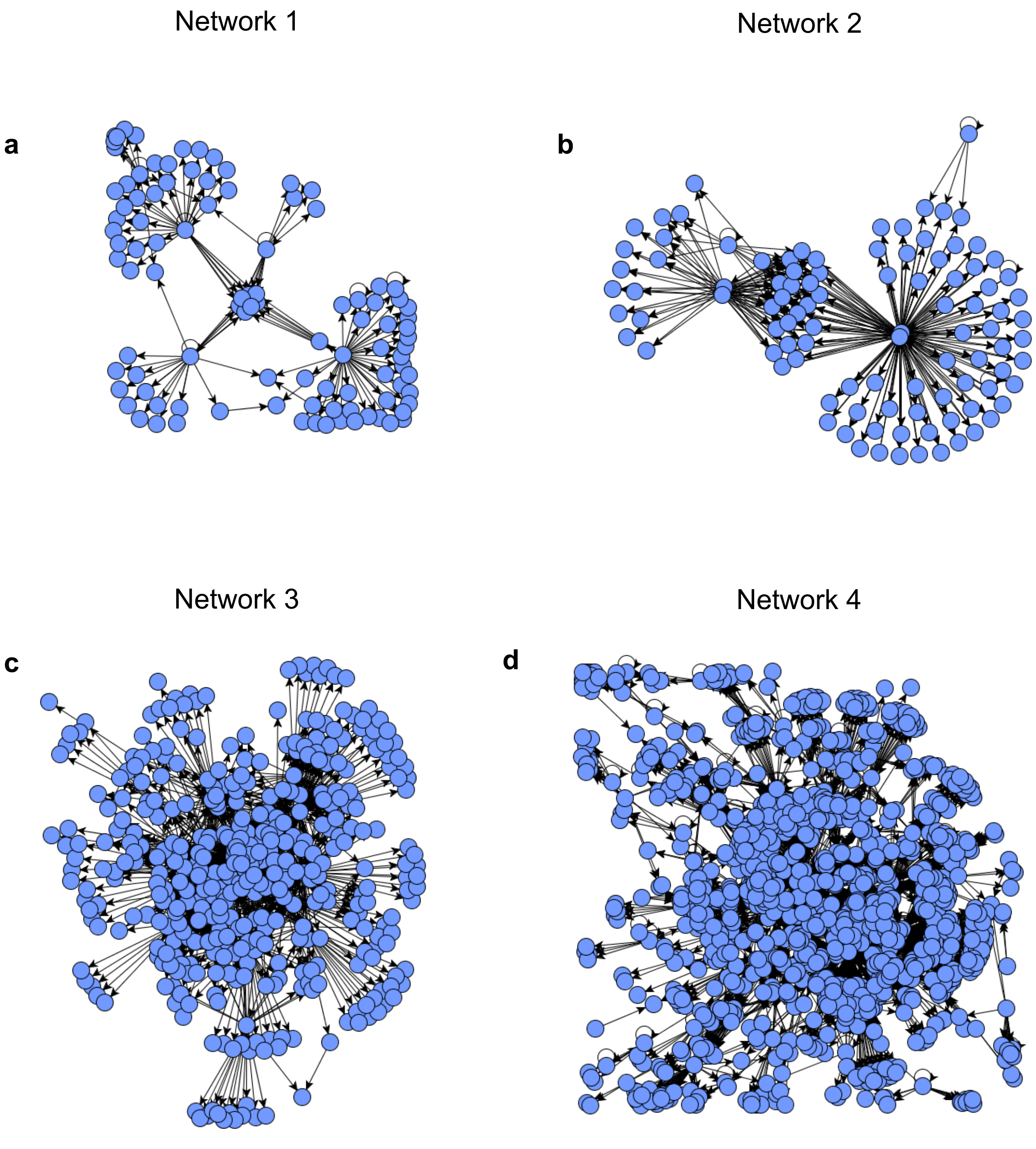


**Figure S1:** The structure of four gene regulatory networks used in this study titled by their network ID. These figures were generated using GNW package[1]. Note that all the auto-regulatory edges as well as cycles were removed prior to feeding networks to Sergio although they are present in this figure. **(a)** Shows network 1, sampled from E.coli, containing 100 genes and 137 regulatory edges. **(b)** Shows network ID 2, sampled from E.coli, containing 100 genes and 258 regulatory edges. **(c)** Shows network ID 3, sampled from S. cerevisiae, containing 400 genes and 1155 regulatory edges. **(d)** Shows network ID 4, sampled from E. coli, containing 1200 genes and 2713 regulatory edges.

#### Supplementary Figure S2

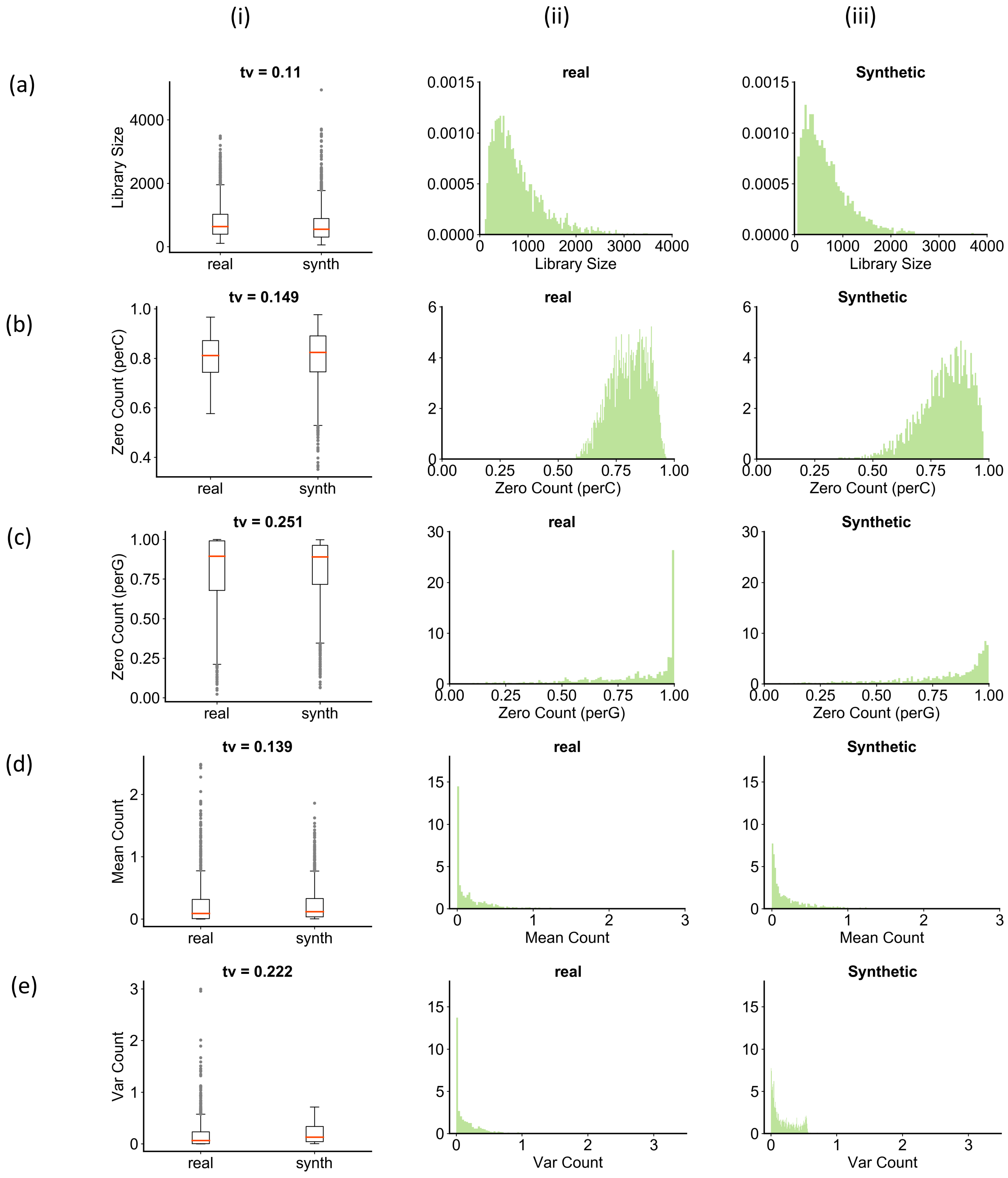


**Figure S2:** To interpret the total variation values and assess the quality of match between the real and synthetic data, for each statistic we looked at a pair of simulated replicate of

DS3 and a real sample which their total variation is close to the median of the total variations of the corresponding statistic. This figure gives a qualitative understanding of the total variation score, which is a number between 0 and 1 reflecting how well two distributions match. Each row represents one of the quantities studied in Figure 2, and shows the distribution of that quantity in one of the simulated replicates and one of the real data sets; the two data sets selected for display here have a total variation ("tv") that is typical for that quantity. **(i)** This column compares the distribution of synthetic against the real data as a box plot. **(ii)** This column shows an alternative visualization of the distribution of the quantity of interest in real data. **(iii)** This column shows an alternative visualization of the distribution of quantity of interest in the synthetic data. The quantities examined include **(a)** library sizes **(b)** zero counts per cell **(c)** zero counts per gene **(d)** mean mRNA counts and **(e)** variance of mRNA counts.

**Supplementary Figure S3**

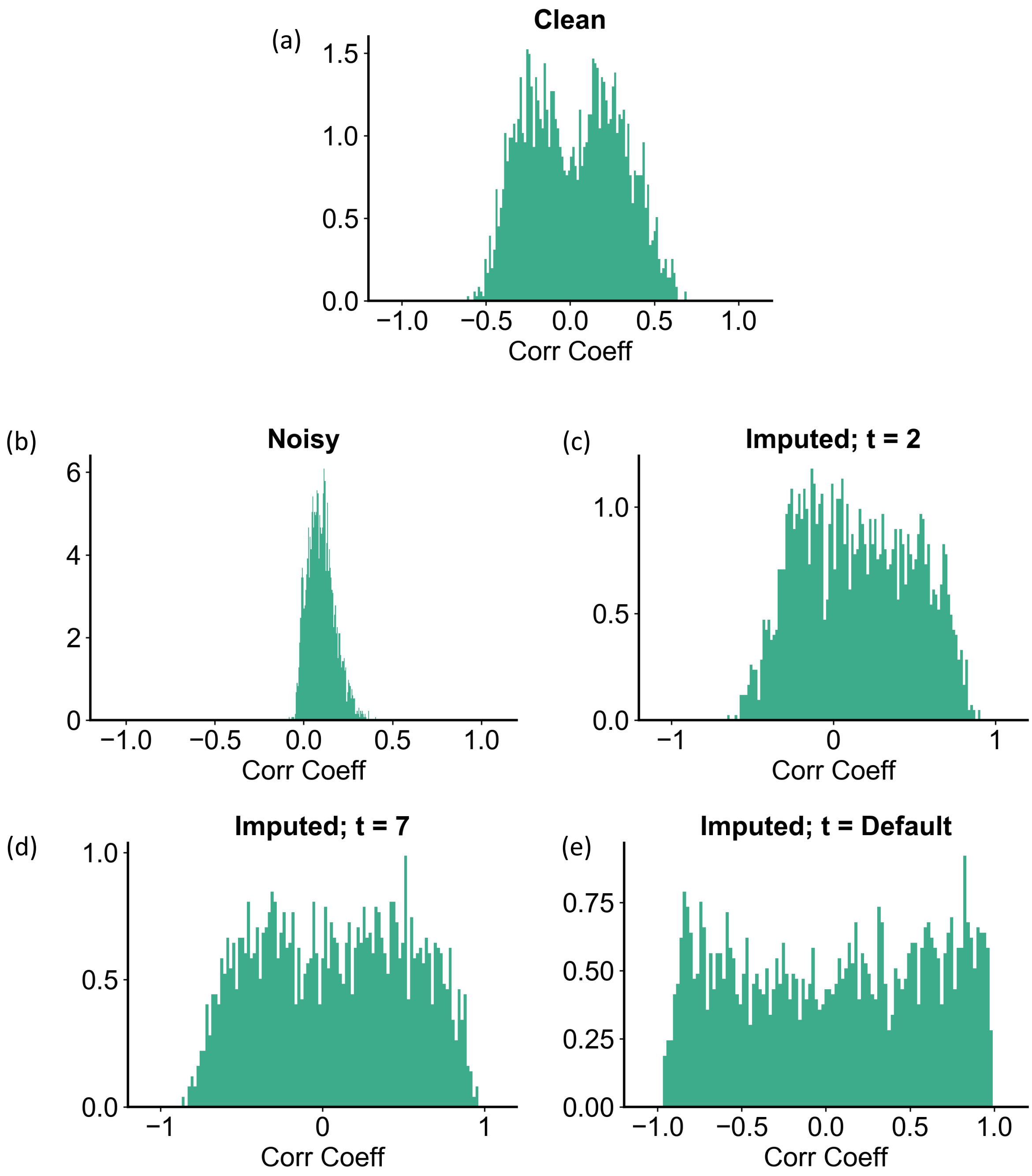


**Figure S3**: Shows the distribution of correlation coefficients between all pairs of interacting genes (regulator – target pairs present in the "ground truth" GRN that was used for simulations) in clean and noisy data of one simulated replicate of DS3, as well as in data imputed by MAGIC[2]. **(a)** Represents the distribution of TF-gene expression correlation coefficients in the clean simulated data. **(b)** Represents correlation coefficients in the noisy data. After adding technical noise, the co-expression signal in the data (panel

a) is severely distorted. **(c)** Distribution of correlation coefficients in the data underlying panel b, after imputation with MAGIC[2] using parameter setting t = 2. Even upon setting *t* to such a small value, several spurious co-expression signals (right tail of distribution as compared to panel a) emerged in the data, compared to the ground truth shown in panel a. **(d)** Distribution of correlation coefficients after imputation with MAGIC using t = 7. This introduces even more false co-expression signals compared to panel c. **(e)** MAGIC imputed data with default *t* setting. We observe almost a uniform distribution over the whole range of correlation coefficients, showing a large number of false positives of co-expressed TF-gene pairs.

#### Supplementary Figure S4

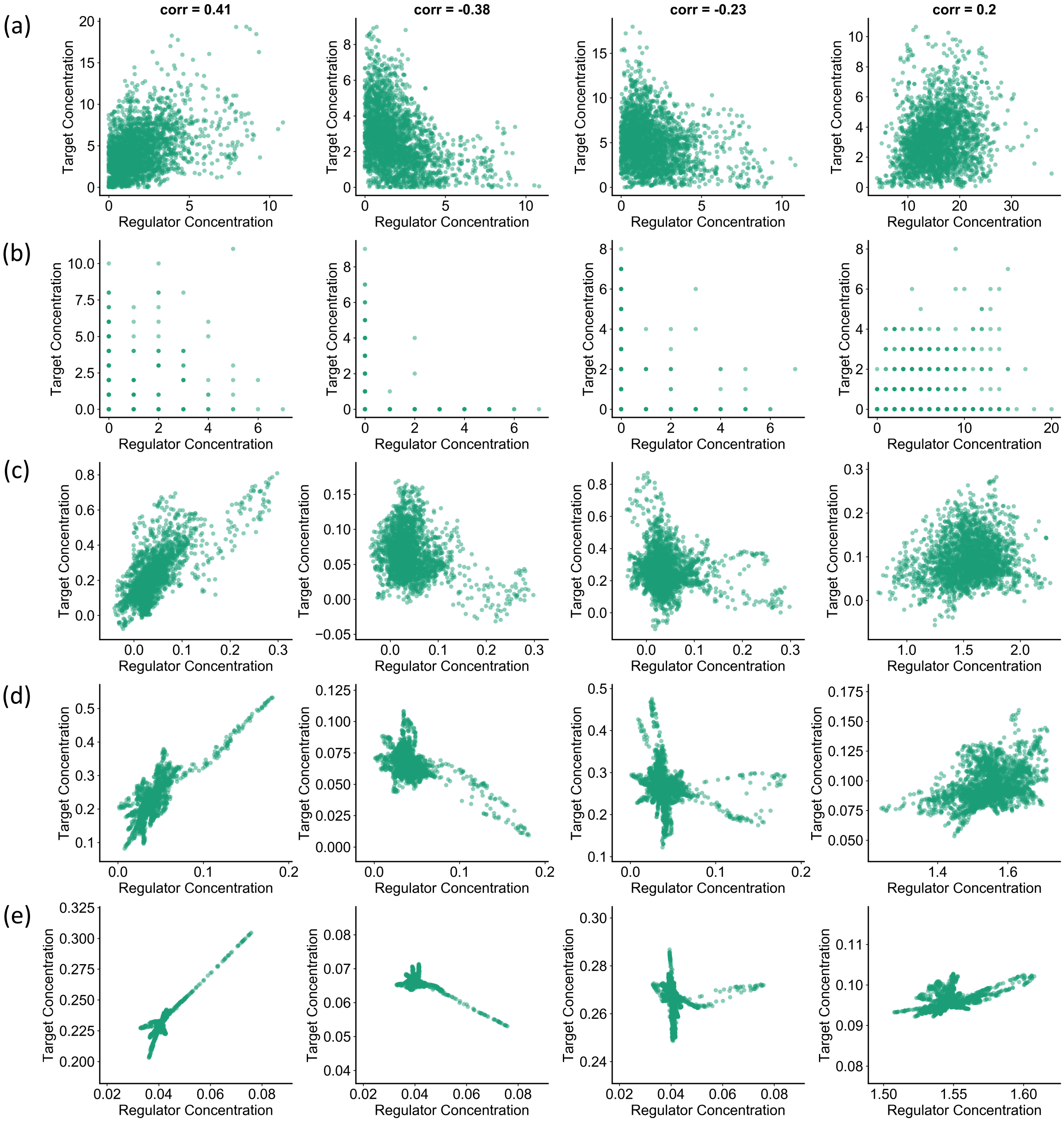


**Figure S4:** Correlation structures in clean and noisy simulated data sets as well as imputed versions of the latter. Columns correspond to four arbitrarily selected regulatory interactions (TF-gene pairs) in DS3 (network 4). **(a)** Clean simulated data. Each panel shows the expressions of the chosen regulator and target pair, in single cells, and the Pearson correlation coefficient between these two observables is noted in caption at the top. **(b)** TF and target gene expression values for the same TF-gene pairs as in (a), after technical noise has been added. The simulated UMI counts are shown. **(c)** TF and target gene expression values for the same TF-gene pairs as in (b), after imputed using MAGIC with $t$ = 2. Note that level of co-expression appears greater than that in clean data ("ground truth"). **(d-e)** Same as (c), but with MAGIC run using $t$ = *7* and $t$ = default respectively.

#### Supplementary Figure S5

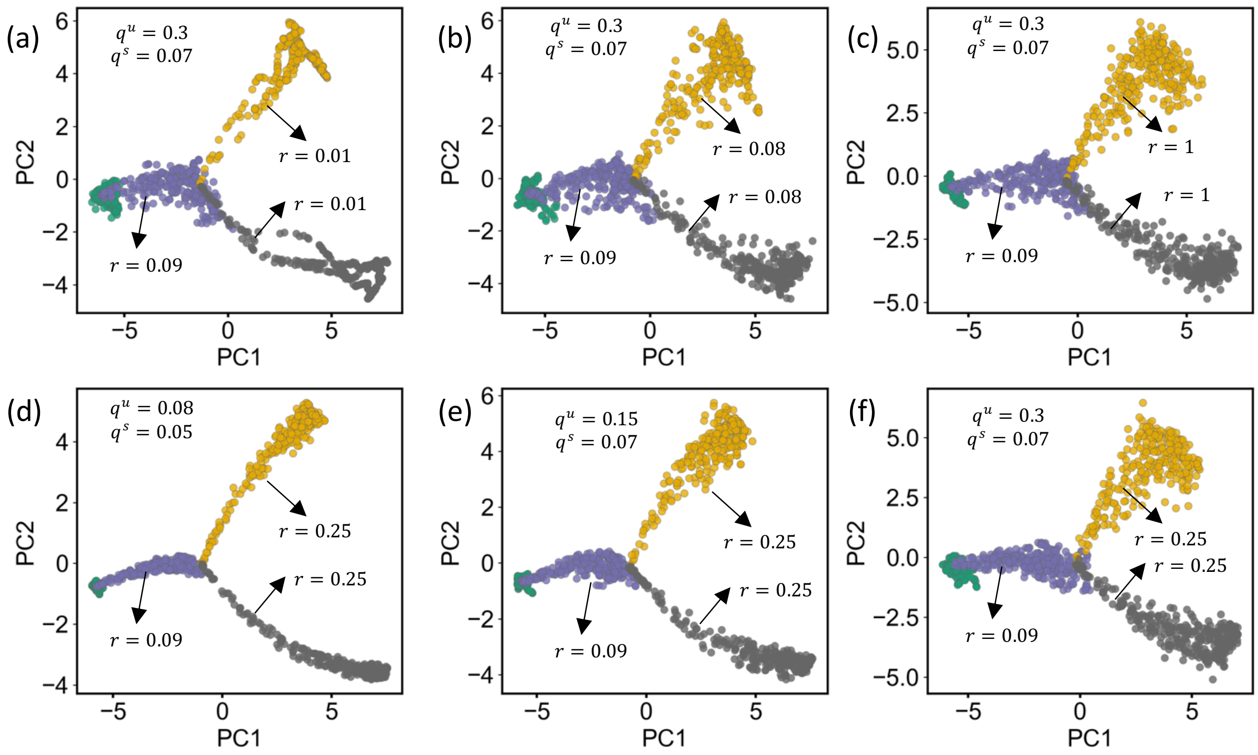


**Figure S5:** User can control the thickness of differentiation path and the dispersion of cells around the trajectory. The user defined migration rate $r$ controls the number of paths that are simulated between two cell types (each edge in the provided differentiation graph). For a given number of cells per cell type ($nCells$) and migration rate $r$, a total number of $r \times nCells$ paths is simulated between the two cell types. Finally, single-cells are randomly sampled from the aggregation of all simulated paths. **(a,b,c)** For fixed unspliced and spliced noise amplitudes $q^u$ and $q^s$ respectively, increasing the migration rate $r$ increases the thickness of the simulated differentiation path as single-cells are sampled from a bigger pool of cells in between the two origin and end cell types. **(d,e,f)** For fixed migration rates $r$, increasing the spliced and unspliced noise amplitudes increases the dispersion of single cells because the higher stochastic noise increases the variance among single-cells.

**Supplementary Figure S6**

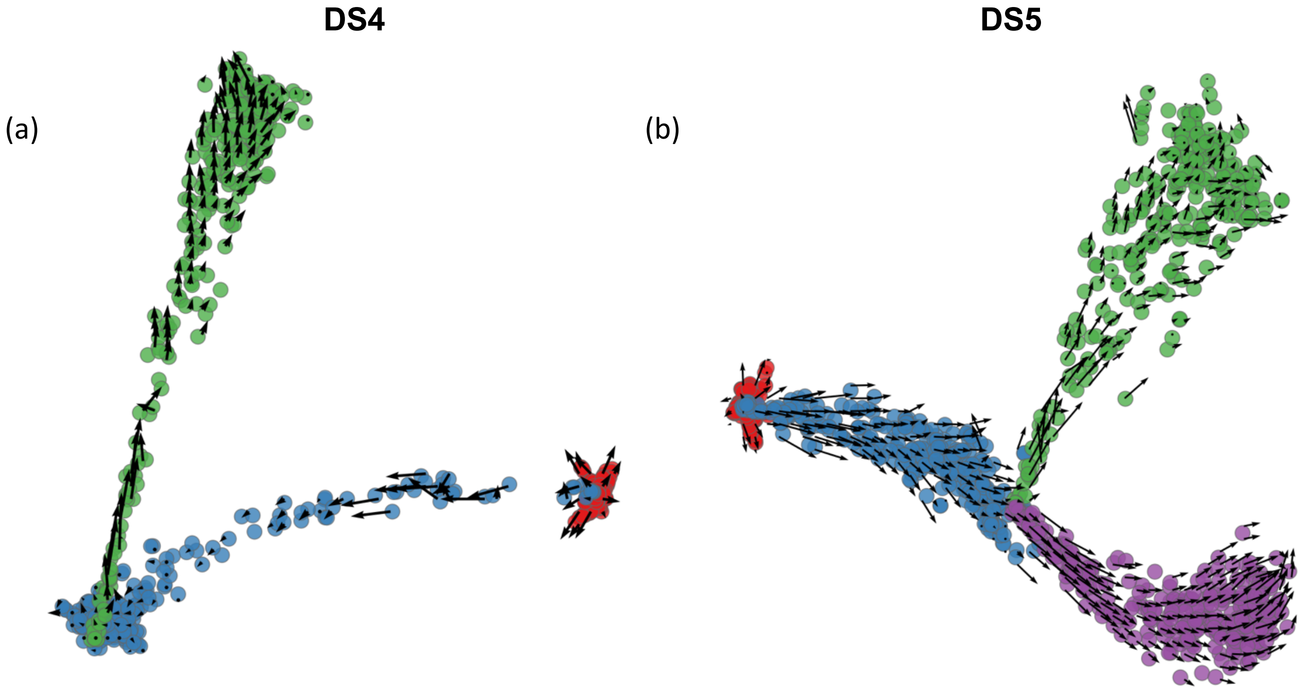


**Figure S6:** Velocity field inferred by Velocyto[3] on clean simulated data in **(a)** DS4 and **(b)** DS5.

#### Supplementary Tables

**Supplementary Table S1:** Technical noise parameters used in this study

| DS-ID | Outlier Genes | | | Library Size | | Dropouts | |
|---|---|---|---|---|---|---|---|
| | $\pi^O$ | $\mu^O$ | $\sigma^O$ | $\mu^L$ | $\sigma^L$ | $k$ | $q$ |
| 1 | 0.01 | 0.8 | 1 | 4.6 | 0.4 | 6.5 | 82 |
| 2 | 0.01 | 0.8 | 1 | 4.6 | 0.4 | 6.5 | 82 |
| 3 | 0.01 | 0.8 | 1 | 4.8 | 0.3 | 20 | 82 |
| 4 | 0.01 | 0.8 | 1 | 4.8 | 0.3 | 20 | 82 |
| 5 | 0.01 | 0.8 | 1 | 6 | 0.4 | 12 | 80 |
| 6 | 0.01 | 0.8 | 1 | 7 | 0.4 | 8 | 80 |

**Supplementary Table S2:** Parameter settings used for running Singe[4]

| Parameter | Set 1 | Set 2 |
|---|---|---|
| $\lambda$ | 0.01 | 0.01 |
| $dT$ | 10 | 5 |
| num_lags | 5 | 9 |
| kernel_width | 2 | 4 |
| prob_zero_removal | 0 | 0.2 |
| prob_remove_samples | 0.2 | 0.1 |
| num_replicates | 2 | 2 |

**Supplementary Table S3:** Low and high expression ranges from which the master regulators' production rates were sampled

| DS-ID | Low Expression Range | High Expression Range |
|---|---|---|
| DS1 | [0.2 0.5] | [0.7 1] |
| DS2 | [0 2] | [2 4] |
| DS3 | [0 2] | [2 4] |
| DS4 | [0 1] | [3 4] |
| DS5 | [0 1] | [3 4] |
| DS6 | [0 1] | [3 4] |
| DS7 | [0 1] | [3 4] |
| DS8 | [0 1] | [3 4] |

### Supplementary Notes

#### Supplementary Note 1: Estimating steady-state concentrations

In steady state-simulations, Sergio approximates the steady-state concentrations of all genes in all cell types prior to starting the simulations. Then, Sergio initializes all the concentrations with their corresponding estimated stead-states values. This is particularly useful for speeding up the simulations since initial concentrations are so close to the values to which the numerical integration is supposed to converge. To do so, Sergio applies a topological sort algorithm on the gene regulatory network in order to layer the graph. After layering the GRN graph using topological sorting, all the regulators of the genes of any layer reside in the preceding layers. Sergio starts estimation of steady-state concentrations (as well as half-response parameters of the hill functions) from the top most layer and continues layer-by-layer until the very last layer of genes. Therefore, all information required for estimating the steady-state concentrations (and half-responses) of genes in the current layer is already available from the user-defined parameters and the estimated concentrations of the genes in the previous layers. Below we demonstrate how Sergio estimates steady-state concentrations.

We start with equation (1) in the Methods section that describes the rate of changes in mRNA concentration $x_i$ of gene *i*:

$$\frac{dx_i}{dt} = P_i(t) - \lambda_i x_i(t) + q_i\left(\sqrt{P_i(t)}\alpha + \sqrt{\lambda_i x_i(t)}\beta\right)$$

In order to reach the steady-state regime, we need to have $\frac{dE[x_i]}{dt} = 0$, where $E[.]$ denotes the expectation operator. Since $\frac{dx_i}{dt}$ is well defined and is bounded, we have $\frac{dE[x_i]}{dt} = E[\frac{dx_i}{dt}]$. So we get:

$$E\left[\frac{dx_i}{dt}\right] = 0 \quad \Rightarrow$$

$$E\left[P_i(t) - \lambda_i x_i(t) + q_i\left(\sqrt{P_i(t)}\alpha + \sqrt{\lambda_i x_i(t)}\beta\right)\right]$$
$$= E[P_i(t)] - \lambda_i E[x_i] + q_i E[\sqrt{P_t(t)}]E[\alpha] + q_i E[\sqrt{\lambda_i x_i(t)}]E[\beta] = 0$$

Also recall that $\alpha$ and $\beta$ are two Gaussian white noise processes which have a zero mean, so we get:

$$E[P_i(t)] - \lambda_i E[x_i] = 0 \quad \Rightarrow \quad E[x_i] = \frac{E[P_i(t)]}{\lambda_i}$$

If gene *i* is a master regulator, according to equation (5) its production rate is solely determined by its user-defined production rate $b_i$ and therefore we can accurately estimate the expected steady-state concertation $x_i$:

$$E[x_i] = \frac{E[b_i]}{\lambda_i} = \frac{b_i}{\lambda_i} \quad ; \quad if\ gene\ i\ is\ Master\ Regulator \tag{S1}$$

However, if gene *i* is not a master regulator, according to equation (5) we get:

$$E[x_i] = \frac{E[P_i]}{\lambda_i} = \frac{\sum_{j \in R_i} E[p_{ij}]}{\lambda_i} \quad ; \quad if\ gene\ i\ is\ not\ Master\ Regulator \tag{S2}$$

where $R_i$ is the set of all regulators of gene *I*, and $p_{ij}$ is a hill function. We use the following approximation for calculating $E[p_{ij}(x_j)]$:

$$E[p_{ij}(x_j)] \approx p_{ij}(E[x_j])$$

Note that this a loose approximation as it clearly over-estimates or under-estimates the true value of $E[p_{ij}(x_j)]$ over a large a range of regulator concertation $x_j$, yet it is good enough for our ultimate goal which is initializing mRNA concentrations. Substituting this back into the equation (S2) we obtain an estimate of the steady-state concertation of gene *i*:

$$E[x_i] = \frac{\sum_{j \in R_i} p_{ij}(E[x_j])}{\lambda_i} \quad ; \quad if\ i\ is\ not\ Master\ Regulator \tag{S3}$$

At the time we calculate the concentration of gene *i* we have already calculated the steady-state concertation of all of its regulators (which reside in the preceding layers of the sorted GRN), hence we have all the information required for this calculation. Note also that the goal of the above estimation is not to find steady-state concentrations *per se,* but to find values close to these concentrations so as to reduce simulation time by starting the simulations at these concentrations.

#### Supplementary Note 2: Controlling the spliced to unspliced count ratio

Employing a similar approach to that discussed in Supplementary Note 1, we can estimate and control the ratio of the expected spliced to unspliced expression in the stationary region of the stochastic transcription process. We start from equation (9):

$$\frac{ds_i}{dt} = \mu_i u_i(t) - \gamma_i s_i(t) + q_i^s(\sqrt{\mu_i u_i(t)}\phi + \sqrt{\gamma_i s_i(t)}\omega)$$

where $\mu_i$ is the splicing rate (decay rate) of the unspliced mRNA and $\gamma_i$ is the degradation rate (decay rate) of the spliced RNA. Recall that $\phi$ and $\omega$ are two zero-mean Gaussian white noise processes. Therefore, under steady-state condition we get:

$$\frac{dE[s_i]}{dt} = E\left[\frac{ds_i}{dt}\right] = 0 \qquad \Rightarrow \qquad \mu_i E[u_i] - \gamma_i[s_i] = 0$$

$$\frac{E[s_i]}{E[u_i]} = \frac{\mu_i}{\gamma_i}$$

Sergio, as an input, takes the decay rate $\mu$ of the unspliced mRNA as well as the splicing ratio $\frac{E[s_i]}{E[u_i]}$; these two together determine the degradation rate of spliced RNA $\gamma$ according to the equation above. This enables the user to simulate gene expressions with any desired spliced to unspliced ratios in order to reproduce different experimental settings.

#### Supplementary Note 3: Adding technical noise to differentiation data

For adding technical noise to differentiation data, we use a similar approach to what we discussed for steady-state data sets. Our implementation supports adding outlier genes, library size effects, and dropouts, before expression values are converted to UMI counts.

**Outlier genes:**

Each gene is designated as an outlier with a user-defined probability. If so, its unspliced as well as spliced expression level (in every cell) is multiplied by a factor sampled from a log-normal distribution, otherwise the expression is left unchanged:

$$\forall i \in \{1 \ldots G\}: \quad \mathbb{I}_i^O \sim Ber(\pi^O) \quad , \quad f_i^O \sim ln\mathcal{N}(\mu^O, \sigma^O)$$

$$\forall c \in \{1 \ldots C\} \quad , \quad \forall i \in \{1 \ldots G\}:$$
$$u_i^c \leftarrow \mathbb{I}_i^O f_i^O u_i^c + (1 - \mathbb{I}_i^O) u_i^c$$
$$s_i^c \leftarrow \mathbb{I}_i^O f_i^O s_i^c + (1 - \mathbb{I}_i^O) s_i^c$$

where *G* and *C* denote the total number of simulated genes and cells respectively, $u_i^c$ and $s_i^c$ denote the simulated unspliced and spliced concentrations respectively of gene *i* in cell *c*. $\mathbb{I}_i^O$ is a binary variable indicating if gene *i* is an outlier, and is sampled from a Bernoulli distribution with parameter $\pi^O$. Also, $\mu^O$ and $\sigma^O$ are user-defined mean and standard deviation of the lognormal distribution from which the outlier scaling factor $f_i^O$ is sampled.

**Library size:**

For every cell (library) a library size parameter is sampled from a user-defined lognormal distribution, and the spliced and unspliced expression of all genes in the cell are scaled such that the total mRNA expression (spliced plus unspliced) of the cell matches the sampled library size:

$$\forall c \in \{1 \ldots C\}: \quad L_c \sim ln\mathcal{N}(\mu^L, \sigma^L)$$

$$\forall i \in \{1 \dots G\} \quad , \quad \forall c \in \{1 \dots C\} :$$
$$u_i^c \leftarrow \frac{L_c}{\sum_{j \in \{1 \dots G\}} u_j^c + s_j^c} u_i^c$$
$$s_i^c \leftarrow \frac{L_c}{\sum_{j \in \{1 \dots G\}} u_j^c + s_j^c} s_i^c$$

where $\mu^L$ and $\sigma^L$ are the user-defined mean and standard deviation of the lognormal distribution from which the library size $L$ is sampled

**Dropouts:**

In real single-cell data it is possible to have expression levels of only the spliced or unspliced RNA of a gene in a cell be zeroed out due to technical noise while the other one is affected by such dropout. To model this, we employ a similar approach as we do for steady-state simulations but add dropout to spliced and unspliced expressions independently:

$$y_0 = q^{th}\ percentile\ of\ Y$$

$$\forall i \in \{1 \dots G\}, \quad \forall c \in \{1 \dots C\}:$$
$$\pi_{i,c}^{D,U} = \frac{1}{1 + \exp\left(-k\left(Y_{i,c}^U - y_0\right)\right)} \quad , \quad \pi_{i,c}^{D,S} = \frac{1}{1 + \exp\left(-k\left(Y_{i,c}^S - y_0\right)\right)}$$
$$\mathbb{I}_{i,c}^{D,U} \sim Ber\left(\pi_{i,c}^{D,U}\right) \quad , \quad \mathbb{I}_{i,c}^{D,S} \sim Ber(\pi_{i,c}^{D,S})$$
$$u_i^c \leftarrow \mathbb{I}_{i,c}^{D,U}\, u_i^c \quad , \quad s_i^c \leftarrow \mathbb{I}_{i,c}^{D,S}\, s_i^c$$

where $k$ and $q$ are two user-defined parameters that determine the logistic probability $\pi^D$, and $Y$ denotes the total mRNA expression matrix in logarithmic scale:

$$Y = \log(X + 1) \quad , \quad X = X^U + X^S$$
$$X^U = \left\{u_i^c; \quad \forall i \in \{1 \dots G\}\ , \forall c \in \{1 \dots C\}\right\}$$
$$X^S = \{s_i^c; \quad \forall i \in \{1 \dots G\}\ , \forall c \in \{1 \dots C\}\}$$

Also we define:

$$Y^U = \log(X^U + 1)$$

$$Y^S = \log(X^S + 1)$$

.

**Conversion to UMI counts:**

Spliced ($UC^S$) and unspliced ($UC^U$) mRNA counts are independently sampled from a Poisson distribution whose mean is the simulated expression level of the gene in the cell:

$$\forall i \in \{1 \dots G\} \quad , \quad \forall c \in \{1 \dots C\} : \qquad UC_{i,c}^U \sim Poisson(u_i^c) \quad , \quad UC_{i,c}^S \sim Poisson(s_i^c)$$

#### References

1. Schaffter, T., Marbach, D. & Floreano, D. GeneNetWeaver: In silico benchmark generation and performance profiling of network inference methods. *Bioinformatics* **27**, 2263–2270 (2011).
2. van Dijk, D. *et al.* Recovering Gene Interactions from Single-Cell Data Using Data Diffusion. *Cell* **174**, 716-729.e27 (2018).
3. La Manno, G. *et al.* RNA velocity of single cells. *Nature* **560**, 494–498 (2018).
4. Deshpande, A., Chu, L.-F., Stewart, R. & Gitter, A. Network Inference with Granger Causality Ensembles on Single-Cell Transcriptomic Data. *bioRxiv* 534834 (2019). doi:10.1101/534834
